# The Inaugural Flatiron Institute Cryo-EM Conformational Heterogeneity Challenge

**DOI:** 10.1101/2025.07.18.665582

**Authors:** Miro A. Astore, Geoffrey Woollard, David Silva-Sánchez, Wenda Zhou, Mykhailo Kopylov, Khanh Dao Duc, Roy R. Lederman, Yilai Li, Yi Zhou, Jing Yuan, Fei Ye, Quanquan Gu, Rémi Vuillemot, Slavica Jonic, Lan Dang, Steven J. Ludtke, Hannah Bridges, Serena Liu, Michael McLean, Valentin Peretroukhin, Johannes Schwab, Eduardo R. Cruz-Chú, Peter Schwander, Marc Aurèle Gilles, Amit Singer, David Herreros, Jose-Maria Carazo, Carlos O.S. Sorzano, J. Ryan Feathers, Ellen D. Zhong, Nikolaus Grigorieff, Pilar Cossio, Sonya M. Hanson

## Abstract

Despite the rise of single particle cryo-electron microscopy (cryo-EM) as a premier method for resolving macromolecular structures at atomic resolution, methods to address molecular heterogeneity in vitrified samples have yet to reach maturity. With an increasing number of new methods to analyze the multitude of heterogeneous states captured in single particle images, a systematic approach to validation in this field is needed. With this motivation, we issued a challenge to the community to analyze two cryo-EM particle image sets of thyroglobulin that exhibit continuous conformational heterogeneity. The first dataset was experimental and the second was generated with a simulator, allowing control over the distribution of molecular structures and enabled direct comparison between participants’ submissions and the ground truth molecular structures and distributions. Participants were asked to submit 80 volumes representing the heterogeneous ensemble and estimate their respective populations in the image sets provided. Participation of the research community in the challenge was strong, with submissions from nearly all developers of heterogeneity methods, resulting in 41 submissions across both datasets. Submissions qualitatively exceeded expectations, with the molecular motions identified by methods resembling both each other and the ground truth motion. However, quantitatively assessing these similarities was a challenge in and of itself. In the process of assessing the submissions, we developed several validation metrics, most of which require reference to the underlying ground truth volumes. However, we have also explored the use of metrics that do not necessarily reference ground truth. This is particularly apt for experimental datasets where ground truth is inaccessible. These approaches allowed us to assess the similarity and accuracy in volume quality, molecular motions, and conformational distribution of di!erent submissions. These metrics and the e!orts of all participants help chart a path forward for the improvements of heterogeneity methods for cryo-EM and for future challenges to validate these new methods as they continue to be developed by the community.

---

Over the last decade, cryo-electron microscopy (cryo-EM) has become mainstream for the determination of high resolution structures of biological macromolecules. Especially for difficult targets, it has become a preferred method over X-ray crystallography and nuclear magnetic resonance [1, 2](Fig. 1A). Several key advantages are that crystals are not a prerequisite, biomolecules can be large, samples can be prepared at near-native environments, and that single molecules are imaged directly. In a typical cryo-EM dataset, images of tens of thousands to millions of biomolecules are captured within a thin layer of vitreous ice. While standard cryo-EM reconstruction methods [3, 4] generally classify these images into a few states so that tens of thousands of images can then be averaged into a single high resolution structure, information about a broader distribution of states can also be captured from cryo-EM datasets due to its single-molecule nature. Indeed, an increasing number of methods are being developed and used to analyze heterogeneity in cryo-EM particles (Fig. 1A), ranging from linear methods to machine-learning algorithms that map the images onto a low-dimensional space representing the main conformational motions, to methods coupled to physical simulators of the conformational motions. We refer readers elsewhere for an overview of approaches [5–7].

**Fig. 1.**
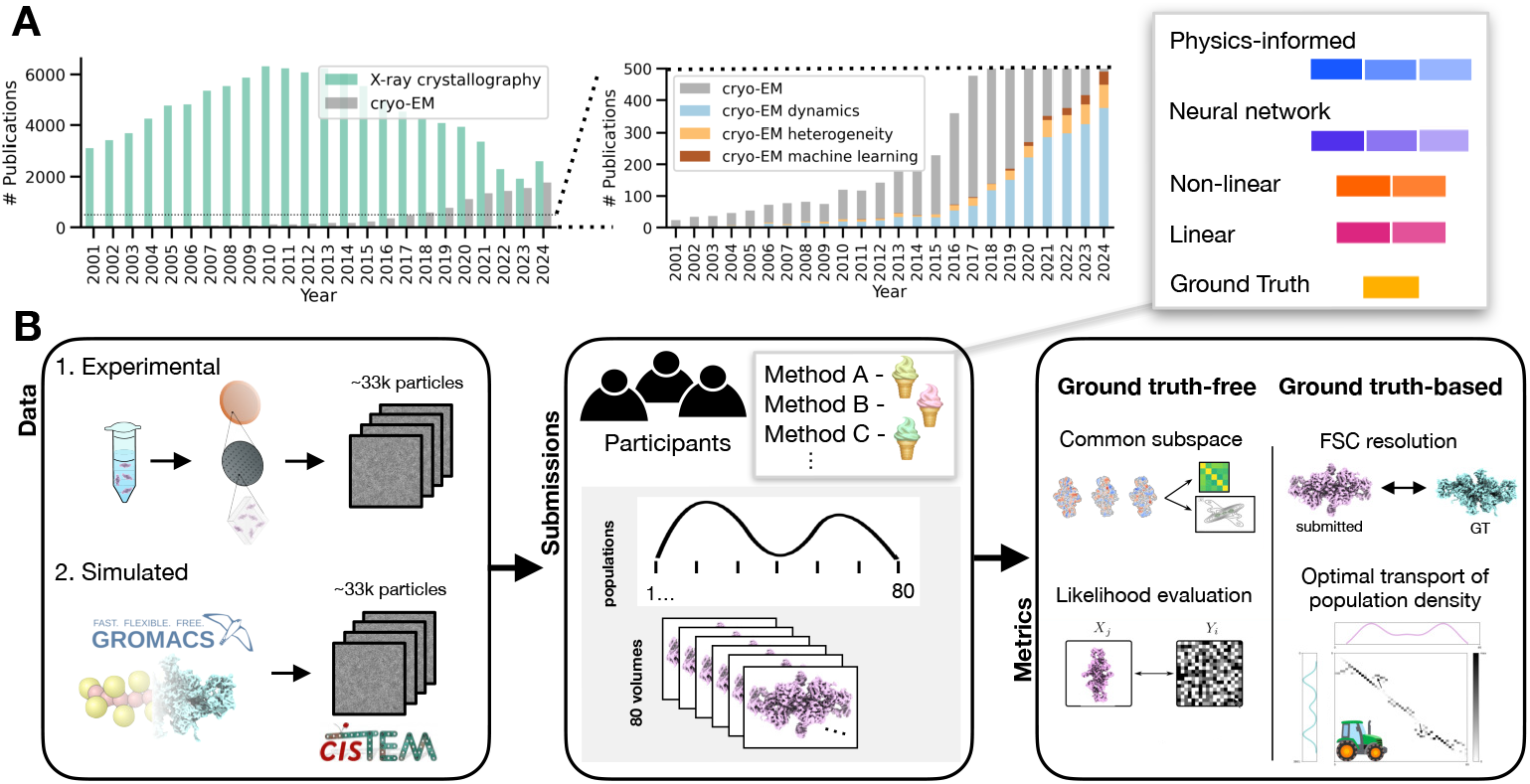
**A**. Number of publications per year for ‘X-ray crystallography’ vs. ‘cryo-EM’ search terms (left) or ‘cryo-EM’ with additional search terms ‘dynamics’, ‘heterogeneity’, or ‘machine learning’ (right). Note zoomed-in y-axis for cryo-EM subterms. Data gathered *via* PubMed Trends. **B**. Schematic of the challenge: dataset generation, submissions, and comparison metrics. Upper right panel indicates the legend for the method type, subsequently used throughout the rest of the manuscript.

The capacity for cryo-EM to capture conformational heterogeneity of biomolecules has potential impacts from basic biology to drug design. Understanding the conformational change, for example, of the SARS-CoV2 spike protein [8, 9] or the capacity for cryo-EM to capture multiple states of drug targets like GPCRs [10] has lead to important therapeutic advancements [11, 12]. Generally, it is acknowledged that static structures of proteins do not tell the whole story of their function. Some key examples of this in cryo-EM heterogeneity analysis include gaining insights into allosteric modulation [13, 14], entropy-driven processes [15], or the transition state of enzymes [16, 17]. However, analyzing the conformational heterogeneity present in cryo-EM data is challenging, because applying different methods to the same dataset can yield varying results [18], and there are limited metrics and tools for quantitatively comparing and validating these outputs [19, 20]. This stems from a fundamental difference in objectives: while traditional cryo-EM reconstruction prioritizes high-resolution structure determination, supported by well-established validation metrics, heterogeneity analysis focuses on resolving conformational motions and estimating population distributions; metrics for validating these aspects remain underdeveloped.

Community challenges in cryo-EM have been launched to compare algorithmic performance on a variety of cryo-EM data processing tasks: CTF estimation [21], volume estimation [22], atomic model building [23], and particle picking [24]. However until now, a community challenge for conformational heterogeneity in cryo-EM has yet to be conducted. One key barrier to running a community challenge for conformational heterogeneity is the lack of metrics for assessing the quality of results. For example, gold-standard metrics that avoid overfitting like the half-set Fourier shell correlation (FSC) for reconstruction or the Q-score for model building do not exist for analyzing output series of volumes from cryo-EM variability methods.

Despite this, the age of machine learning has made it clear that benchmarks and metrics can rapidly accelerate algorithmic advances [25], especially obvious in the field of structural biology with the role of CASP in facilitating the success of AlphaFold [26]. Motivated both by these successes, and our own questions about how much to trust the results produced by conformational heterogeneity methods in cryo-EM, we launched the Flatiron Institute Cryo-EM Conformational Heterogeneity Challenge to the community in the summer of 2023, which included two datasets: one simulated with a ground truth, for which metrics would be more straightforward, and one experimental dataset. We had strong participation from the cryo-EM method’s developer community across the globe, with 41 submissions for both datasets. Here, we first describe the construction of the challenge, and the participant’s submissions. Then, we present the metrics we developed to assess and compare the methods’ outputs for several specific tasks, assessing volumes, motions, and distribution quality across submissions. Overall, we do not find a single method that consistently outperforms the others; rather, different methods highlight different trends.

## 1. Results

### 1.1 Dataset design

In the design of this heterogeneity challenge, we sought a system that, though simple, would present a non-trivial task for the participants. We were also motivated to include both experimental and simulated cryo-EM images (Fig. 1B, left) that exhibited continuous conformational heterogeneity. The experimental dataset was a 33,742 particle stack of thyroglobulin that reconstructed to a resolution of 3.1 Å (see Methods for details). We subsequently used this experimental dataset to design the simulated dataset. The simulated dataset was constructed using the Euler angles, defocus values, and particle number of the experimental dataset, and was generated using the *cis*TEM (Computational Imaging System for Transmission Electron Microscopy) simulator [27, 28]. This simulator is more complex than simulators that only make the typical weak phase and projection approximations and employ Gaussian white noise.

We chose this more complex simulator because we wanted to model experimental cryoEM data more closely with *cis*TEM’s multislice integration, explicit solvent modeling, and inclusion of radiation damage. Notably, not all participants were able to distinguish between the experimental and simulated datasets during the initial phase of the challenge because the simulated dataset exhibits a higher level of noise compared to the experimental dataset..

A main motivation for including a simulated dataset in this challenge was to have ground truth conformations and a distribution to which we could compare participants’ submissions. We ran coarse-grained molecular dynamics simulations of thyroglobulin whose conformational space we then redistributed along the first principal component of its motion in molecular space (MD-PC1), which we defined as the “ground truth heterogeneity coordinate” (see section 3.1). We re-weighted the conformational distribution according to a nonuniform mixture of three Gaussians that included a less-populated state in the middle of this space, which we believed would represent an interesting challenge for methods to identify (Fig. 4). A total of 3,861 conformations were sampled from this distribution, backmapped to atomic coordinates and used to generate the 33,742 image simulated particle stack. While re-weighting the distribution along MD-PC1 and quenching motions MD-PC2 (see Methods for details) was done to emphasize one degree of freedom, the simulated volumes still contained other degrees of freedom and exhibit considerable side chain motion.

### 1.2 Submission summary

For both of the provided datasets, we asked for participants to submit 80 volumes along the dominant degree of freedom and the relative populations of each volume (Fig. 1B, middle). This format was influenced by our knowledge that most cryo-EM heterogeneity methods produce volumes as output, rather than molecular ensembles for example, and the request for 80 volumes was to increase the chance of picking up low-probability states. To facilitate comparison, we asked that participants align their submitted volumes to a provided reference and match its voxel size (2.146 Å). Additionally, we designed two mock submissions for the simulated dataset built from the Ground Truth (GT) which we refer to as Averaged GT and Sampled GT. To generate these, first we sorted the volumes by MD-PC1, and split up the volumes into 80 equispaced groups with an approximately equal number of volumes. Averaged GT consists of 80 volumes generated by Euclidean averaging each equispaced set. Sampled GT consists of 80 volumes selected by choosing the most commonly sampled volume in each equispaced set.

All submissions underwent the same pre-processing of being aligned to the consensus reconstruction for experimental dataset, or to the average of all the ground truth volumes for the simulated data, before being passed through the separate analysis pipelines (see Methods for further details). This was done to avoid favoring one volume pair, or having to perform multiple alignments between all pairs. We labeled the submissions by ice cream flavor to preserve the anonymity of the participants. We have grouped methods, as shown in the top right of Fig. 1, as physics-informed (blue), neural network without physics (purple), non-linear (orange), linear (magenta), and ground truth for the simulated datasets (yellow). We note that some ice cream flavors have additional number labels (e.g. Ice Cream Flavor 1, Ice Cream Flavor 2, etc). These count the number of rounds the same method made a submission. Note that only submissions from “round 1” are blind; after that participants had seen a presentation illustrating the ground truth distribution for the simulated dataset (although not the ground truth latent values of the simulated images, and not the ground truth volumes).

### 1.3 High level overview of submissions

As an initial analysis of the submissions, the simplest procedure is to visually inspect the submitted volumes and the conformational motions (Fig. 2A). From a qualitative perspective, the submissions were diverse in terms of resolution and, in general, the motions represented were consistent (Supplementary Movie 1). Despite asking for submissions to be aligned to a reference volume at a specific pixel and box size, substantial pre-processing of the submitted volumes was still required. For the simulated dataset, since the ground truth is available, we could also assess the resolution of the submissions by comparing the volumes to the ground truth structures using the Fourier shell correlation (FSC) with a 0.5 threshold (Fig. 2E). We computed the FSC for all 3861 GT volumes against all volumes in every submission, and matched each of the 80 volumes with the closest GT volume using the resolution at FSC 0.5 threshold. Some submissions had relatively tight resolution distributions, either higher (Cookie Dough 1 around 9 Å; Mango 1 around 9.5 Å) or lower (Chocolate Chip 1 and Rocky Road 2 around 18.5 Å). Other submissions had a large range of values, for example Chocolate Chip 2, Salted Caramel 3. As an alternate validation metric not needing access to ground truth volumes, we directly assessed each submitted volume against a subset of the individual images by exhaustively searching the pose space and identifying the optimal cross-correlation for each image (see Methods for details). We observed that the distributions of image-to-volume cross-correlations are broad for both the experimental and simulated datasets (Fig. 2C and F, respectively). However, some methods produce volumes that align more closely with the images than others. Notably, the patterns observed in image-to-volume comparisons do not always align with those seen in volume-to-volume evaluations.

**Fig. 2.**
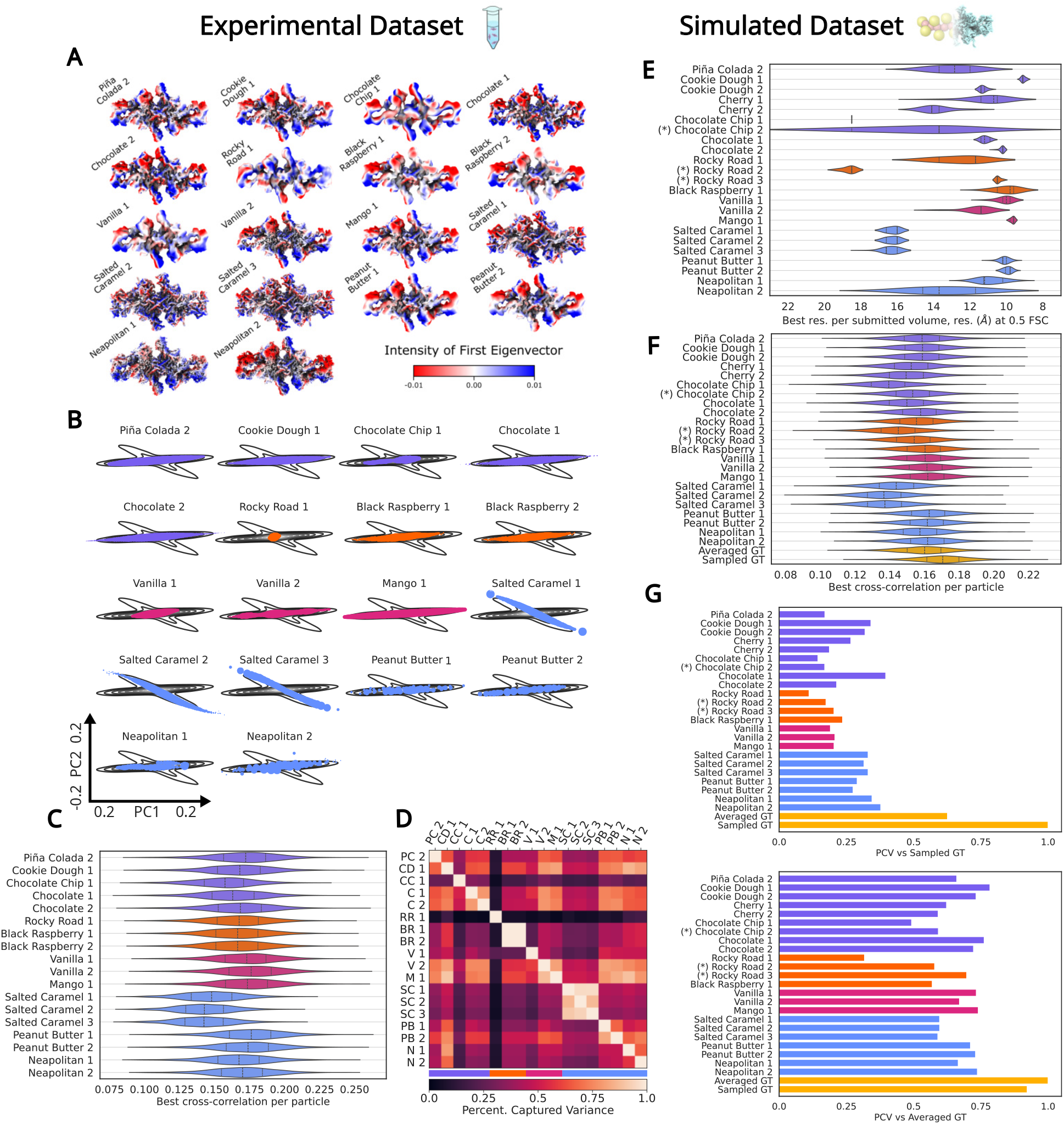
**A**. The 40th volume from each submission for the experimental dataset, colored by the first PCA eigenvector (blue is positive, red is negative, white is zero). **B**. Projection of submissions onto PC1 and PC2 of the common subspace for the experimental dataset. Each dot represents a submitted volume, with the size defined by the submitted population. The contour plot is a weighted kernel density estimate computed with the projection of all submissions and their populations. In **C**. and **F**. the best volume-to-image cross-correlation for each particle is shown for the experimental and simulated dataset, respectively, for a subset of particles. **D**. The percentage of captured variance (PCV) between the submissions in the experimental dataset, specifically, the amount of variance captured between each pair of submissions. Similarly, G. shows the PCV between the submissions and the ground truth. E. Best resolution for each submitted volume obtained by computing the Fourier shell correlation (FSC) against the GT volumes. The resolution is defined using the 0.5 FSC threshold. (*) Indicates submissions that made use of an expanded simulated dataset with 674,840 particles.

### 1.4 Comparing conformational motions through PCA

For both experimental and simulated datasets, a subspace can be computed for each submission. These subspaces can then be quantitatively compared, and a common subspace can be derived to visualize the similarity between the inferred conformational motions. This is accomplished through Principal Component Analysis (PCA) on each submission, which provides an orthogonal basis defined by the eigenvectors V and the eigenvalues *S*^2^ of each submission’s volume covariance matrix. These bases can be used to compute a common subspace, as detailed in Section 6.1.4. We compute the similarity between the different bases using the Percentage of Captured Variance (PCV) introduced in ref. [29].

Fig. 2A-B shows the behavior of each submission in both the individual and common subspaces for the experimental dataset. Fig. 2A illustrates the regions of variance in the first principal component eigenvector for each submission’s embedding. A change in color indicates regions of variance between the submitted volumes, providing insights into the motions predicted by each submission. Conversely, Fig. 2B shows the projection onto PC1 and PC2 of the common subspace obtained for the simulated dataset submissions. Most methods cover a similar range in PC1, demonstrating that most methods share at least one high variance conformational motion. The dominance of Salted Caramel in the second principal component can be attributed to it having more submissions and possibly presenting higher frequency variance. Supplementary Fig. S1B shows the projection of the motions for the simulated dataset, and the conformational motion with the highest variance aligns well with that of the ground truth and most of the submissions.

To quantify the similarity between different submissions, we measure the PCV between each pair of submissions, which we denote by *PCV* (*i, j*), where *i* and *j* refer to the *i*-th and *j*-th submission, respectively. Submission *i* serves as the reference, and we compute how much of its variance is captured by the subspace of submission *j* (see Methods for details). The asymmetry in this metric allows us to quantify, when comparing two submissions, which submission is more e”ective at capturing variance. For instance, when comparing the two ground truth mock submissions through this metric in Fig.2G, we observe that the Sampled GT e”ectively captures the Averaged GT, while the opposite is not as true. This finding aligns with our expectations, as averaging tends to result in the loss of high-frequency features. Fig. 2D illustrates the variance captured among the submissions in the experimental dataset. In this figure, the rows correspond to the reference submissions (i.e., submission *i*). We notice that submissions produced using the same method, as well as those belonging to the same category, tend to cluster together. This is evident from the higher PCV values that are situated close to the diagonal.

Fig. 2G shows the PCV for the simulated dataset using the Ground Truth mock submissions–Averaged GT and Sampled GT–as references, top and bottom, respectively. The results indicate that methods with higher resolution outputs (e.g., Salted Caramel, Chocolate, and Neapolitan) are better at capturing the high frequency features of the conformational heterogeneity from the ground truth volumes. Once these features are averaged out, lower resolution submissions (e.g., Rocky Road, Vanilla, Mango) perform much better. There are also methods that have a good balance of both cases, such as Chocolate and Cookie Dough. Supplementary Fig. S1C shows the percentage of captured variance between all methods as a distance matrix. As for the simulated dataset (Fig. 2D), we observe that methods assigned to the same categories have a tendency to form clusters, which can be seen from bright spots along the diagonal. Additionally, we notice that Piña Colada, Peanut Butter and Chocolate, methods that presented a balanced behavior in the simulated dataset, are easily captured by other methods (bright rows); while Neapolitan and Peanut Butter are good at capturing other methods (bright columns).

### 1.5 Comparing conformational distributions

A key objective of the challenge was to assess the capacity for heterogeneity methods to predict the population distribution along the 1D conformational coordinate. Since the true populations are unknown for the experimental dataset, our analyses will first focus on the simulated dataset, where the ground truth distribution is available.

#### 1.5.1 *Via* linear subspace embedding

One way to compare the conformational distributions is by using the linear subspaces introduced in Section 1.4. Specifically, we projected the submissions onto the subspace defined by the Sampled GT and visualized the distributions with respect to the first principal component along with the population of submitted volumes. This is illustrated in Supplementary Fig. S2, where these distributions are compared to the distribution of the Averaged GT. Most submissions report a distribution with two modes, either missing the middle mode or showing shifted modes that possibly accumulate the middle mode into the lateral modes. Other submissions simply report a single mode in the middle. Only three submissions (Cherry 2, Chocolate 2, and Vanilla 2) infer the three modes present in the ground truth, although corresponding to the second (i.e. non-blind) round of submissions. On the other hand, most submissions obtain a good coverage of the first principal component, showing that most methods are good at estimating the conformational variability coverage.

#### 1.5.2 *Via* Earth Mover’s distance and Kullback–Leibler divergence

In order to quantitatively compare the information in the volumes and their population weights, we used a distance over distributions known as the earth mover’s distance (EMD) [30]. The EMD compares two probability distributions over a metric space and is also known as the “Wasserstein distance” or “Kantorovich–Rubinstein metric” in the optimal transport literature. In discrete settings it involves finding a soft correspondence matrix between two weighted spaces, referred to as an optimal transport plan, which is a joint probability matrix between these spaces. In other words, the optimal transport plan is the joint distribution that minimizes the distance, *d*, defined between these spaces. Here we compare two discrete probability distributions (of size 3861 for the ground truth, and 80 for the submission) where we use the EMD, or alternatively the Kullback-Leibler (KL) divergence, with a volume-to-volume distance, *d* (see Section 6.2 for details). The EMD has already been used to compare distributions of atomic models of biomolecules [31], but in our case, we compared voxelized volumes, which do not generally include correspondence information between labeled coordinates.

We experimented with various “volume-to-volume” distance functions suitable for comparing these discretized scalar densities (broadly BioEM3D, FSC, Zernike3D, transport distances — see Methods for details), balancing computational feasibility with robustness to pose and high frequency conformational heterogeneity. Furthermore, since inferring the relative populations of their submitted volumes was a new task for many challenge participants, we also optimized their relative populations under the EMD and KL objectives. In other words we used (i) ground truth volumes and (ii) ground truth relative populations, and (iii) submitted volumes, but optimized the submitted relative populations.

As reference, we present here the results using one specific volume-to-volume distance, the BioEM3D distance, which marginalizes out the overall o”set, magnitude and noise level of the submitted volumes, under a Gaussian white noise model in 3D voxel space. Results for other volume-to-volume distances are shown in supplementary Section 6.4. In Fig. 3A, we show the volume-to-volume distance matrix for several representative submissions. Using these, we compare the optimal relative population (colored) to the submitted one (black) in the submission’s volume space (Supplementary Fig. 3B). As illustrated by the Averaged GT, the optimized populations match the reference with high accuracy. Also note that with the BioEM3D distance in an EMD volume distribution objective yielded a smooth optimal population for Sampled GT, which was also seen for the KL conformational distribution objective, but not seen in the other volume-to-volume distances (data not shown).

**Fig. 3.**
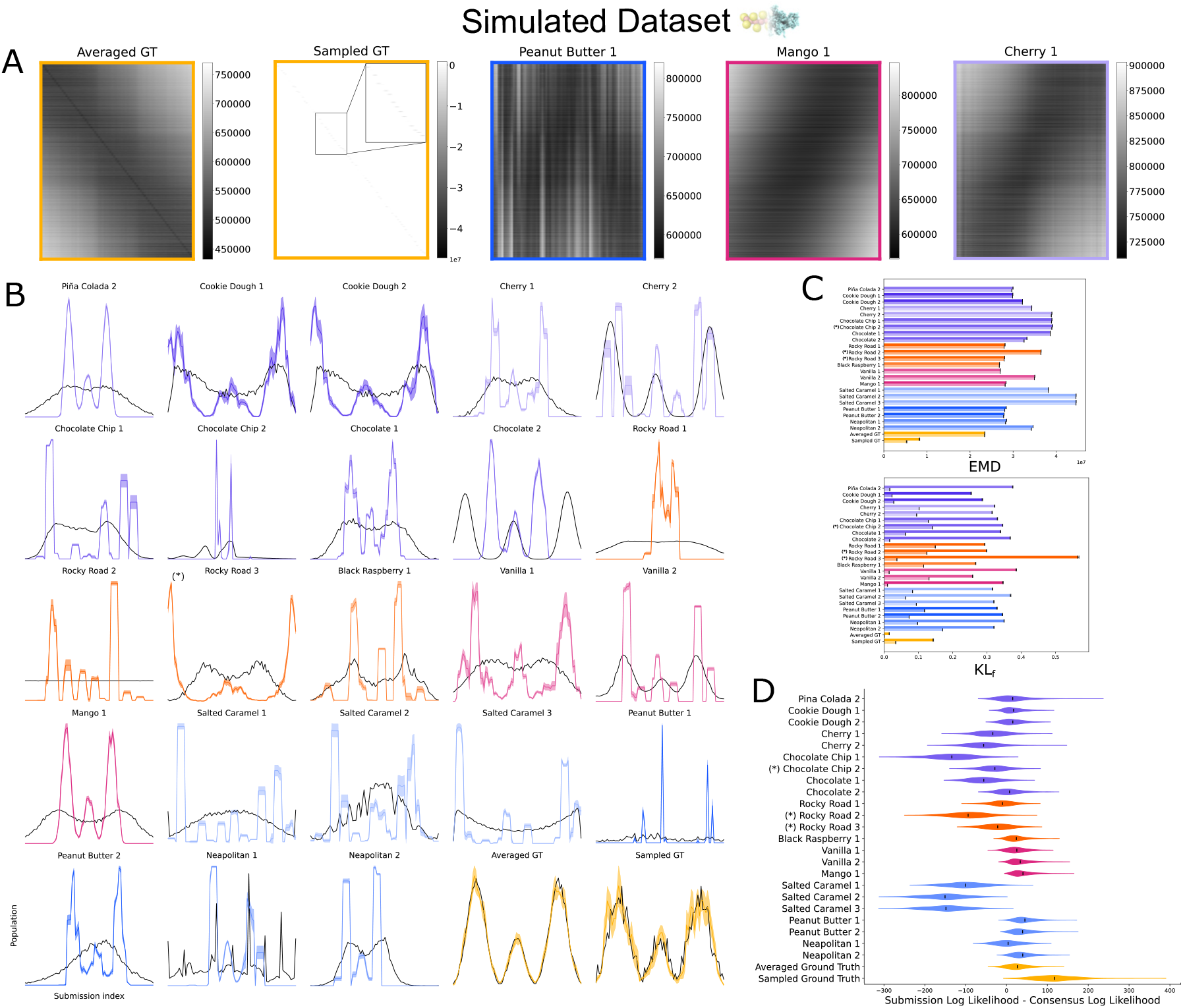
**A**. Volume-to-volume BioEM3D distance, Eq. (17), for select submissions. The distance matrices for all submissions can be found in Supplementary Fig. S3. **B**. Optimal population weight (in color, Eq. (33)) *via* the earth movers distance (EMD), and original submitted population weight (black). Replicates average volume-to-volume distances over 50 nearest neighbors by ground truth ordering (MD-PC1). **C**. EMD and KL_*f*_ comparing submitted volumes and population weights to the ground truth. The error bars are from averaging using different neighborhood sizes of 40,45 and 50. **D**. Ensemble-to-image likelihood (Eq. 39) for a subset of 1224 images corresponding to Sampled GT, relative to the volume-to-image likelihood of the consensus GT volume.

We averaged the volume-to-volume distance matrix along neighborhoods of ground truth ordering (MD-PC1). However, we observed that when large neighborhoods are used and the number of (averaged) ground truth states subsequently decreases, the sparsity of the solution increases. This is expected with the unregularized EMD and KL objectives. We therefore averaged over random replicates, where each replicate drops 90% of ground truth volumes. While each replicate yields sparse optimized relative populations, the average is more smooth. In order to promote smoothness without assuming a ground truth ordering we used entropic regularization, but it did not give the same trend (Supplementary Fig. S4).

Supplementary Fig. S5D compares the conformational distribution with the EMD using a BioEM3D volume-to-volume distance, and different trends are seen even among submissions of the same ice cream flavor: for instance Cherry, Chocolate, Rocky Road, Vanilla, Peanut Butter, Neapolitan, and finally Sampled/Averaged GT. Note that while almost identical results are seen for the weighted populations optimized by a KL vs. EMD objective (data not shown), the magnitude of change in the KL objective is more pronounced (Fig. S5C). Small changes in the EMD objective are expected since the volume-to-volume distances have an intrinsic scale: for example, Averaged GT ranges from 0.4 - 0.75 (Fig. S5A).

In order to put submissions on the same scale, we rank normalized the volume-to-volume distances. Supplementary Fig. S6 shows that rank normalization had a minimal e”ect on the optimized relative population (although Sampled GT did dramatically change), but changed the trends of the objective values. Note that the BioEM3D volume-to-volume distance for Sampled GT atypically shows low values on the diagonal, and is high elsewhere, regardless of the proximity of the submitted volume along the ground truth ordering (MD-PC1), which could explain why its behavior was different to other submissions. This e”ect is also seen in the BioEM3D volume-to-volume distance matrix of Averaged GT compared with itself in Supplementary Fig, S4B (lower). In both cases there is an exact match between some volume pairs being compared, which have a BioEM3D value several order of magnitude lower than non-exact matches.

For roughly half of the submissions, Fig. 3B illustrates that the optimized probabilities are a trimodal mixture, with two symmetric large modes close to the two large modes from the submitted population and another mode ranging from around half to a third its size in the middle, which resembles the ground truth distribution, as expected. In a related way, various bimodal submissions are optimized to be trimodal, indicating the difficulty of inferring low probability surrounding a “hidden” middle mode. This is not surprising given the high level of noise in the images, small scale of the motion, and low number of images that represent the trough in the ground truth relative population distribution.

### 1.6 Ensemble-to-images comparison metric

The previous metrics for assessing distributions relied on ground truth volumes and populations, making them infeasible in the evaluation of submissions for the experimental dataset. Here, we introduce a comparison metric that bypasses this requirement by directly evaluating how well an ensemble (i.e., volumes and populations) explains a given set of cryo-EM images. This metric is independent of identifying a common subspace, and does not require the use of ground truth volumes, because it directly compares volumes with populations to the images using the ensemble-to-image likelihood (Eq. 39) [32]. Each volume is compared to each image by finely searching the pose and particle center, assuming a forward model of image formation with Gaussian white noise [33]. As this pose-search is computationally expensive, only a subset of particles from each dataset were chosen to evaluate this metric (see Section 6.6 for details). The higher the log-likelihood the more likely the particle images are to be generated by the volumes at the given relative population. With this metric, we ranked the performance of submissions for 2000 of the experimental images (Supplementary Fig. S7), where we had no ground truth for the poses or volumes. While for the simulated dataset, we specifically chose to compare to a subset of 1224 images which corresponded to the 80 volumes in the Sampled GT, allowing us access to a ground truth comparison for this metric (Fig. 3D). These results show that the log-likelihoods from the ensemble-image comparison agree to some extent with the EMD/KL results, especially for the submissions that are far from the data (e.g. Salted Caramel, Rocky Road 2, Chocolate Chip Unlike the EMD/KL, many submissions show better agreement with the images than Averaged GT. However, the volumes corresponding to the exact images (Sampled GT) have the best agreement. In this case, we found we could use Markov chain Monte Carlo (MCMC) and ensemble reweighting [32] to recover the correct relative populations between the volumes in the Sampled GT submission directly using the image-to-structure likelihoods (Supplementary Fig. S8). Overall, this suggests that the ensemble-to-image likelihood comparison is able to discern if the ensemble is a poor match, but is sensitive to an exact match between the images and volumes, and care must be taken if volumes are smoothed or averaged in some manner.

When comparing the log likelihood results for submissions to both the experimental and simulated datasets, we find that no submissions perform significantly better or worse on either dataset. Similar trends between the relative ranks of submissions are observed in Fig. 3D and Supplementary Fig. S7. This can also be seen in the cross-correlation scores (Fig. 2C, F). However, because of higher noise levels in the simulated dataset, the difference between the log likelihoods compared to the consensus structure in the submissions to the simulated dataset are usually less than those for the experimental dataset.

## Discussion and conclusions

The capacity to accurately discern the structures and populations of conformational ensembles of biological macromolecules has vast implications for the understanding of biological processes. Cryo-EM o”ers an unprecedented window into this conformational heterogeneity. However, cryo-EM methodologies and validation metrics have primarily focused on achieving high-resolution reconstructions, while quantitative frameworks for assessing the accuracy of approaches that recover conformational motions and population distributions are still emerging. Here, we presented both experimental and simulated datasets analyzed by different methods in the hands of the developers themselves and subsequently compared to one another and to ground truth when available. The submissions overall were able to capture similar conformational motions and many methods even captured populations comparable to the ground truth distribution in the simulated dataset.

We launched this challenge knowing that the metrics with which the submissions would be assessed had yet to be developed. Therefore, in the process of analyzing the challenge submissions, we have developed and tested new quantitative cryo-EM heterogeneity metrics, with the aim of providing computationally tractable, reliable indicators of motions and population prediction. We developed three distinct classes of metrics to evaluate the quality of submissions across different tasks: *i* ) a PCA-based method that compares conformational subspaces, *ii* ) an optimal transport method that allows for the comparison of populations under volume-to-volume similarity measures, and *iii* ) an image-to-ensemble likelihood that allows for direct comparisons of the volumes and populations to the provided images. One common advantage of all these metrics is that they do not require analysis of the latent spaces produced by the heterogeneity analysis methods themselves. One issue we have had with all metrics, however, is their sensitivity to the resolution of the submitted volumes and other aspects of the preprocessing of the volumes. When comparing volume series through PCA, it is important that the volumes in all series are close in resolution, as we found that otherwise methods with high resolution outputs would be unfairly penalized. We solved this issue by normalizing the power spectrum of submissions and ground truth volumes, allowing us to find that submissions like Neapolitan, Salted Caramel, and Peanut Butter excel at matching the ground truth volumes. For the volume-to-volume metrics, one can downsample volumes or smooth out the distance to high frequency information (e.g. truncate the FSC at a lower resolution). For the volume-to-images comparison, one can likewise simply downsample images and volumes.

Overall, while some methods show similar patterns, no single method consistently outperforms the others; instead, different approaches demonstrate strengths in different areas. We also did not observe strong evidence that particular group types systematically outperform others across specific tasks. For instance, some ML-based methods—such as Cookie Dough—achieved strong EMD scores but performed average in terms of image cross-correlation. Likewise, performance among physics-based methods varied significantly depending on whether the metric related to resolution. These findings suggest that the specific design and implementation details of each method play a critical role in determining its e”ectiveness for a given task.

Throughout the analysis of the challenge submissions, some of the limitations of the dataset choice and submission format have become clearer, though ultimately we believe this task has been a suitable one for a first attempt at a conformational heterogeneity challenge in cryo-EM. For example, while thyroglobulin is a biomolecule of typical size (660 kDa) and conformational heterogeneity can be detected with a modest particle count (33,742 images), it is a symmetric molecule, hence conformational heterogeneity can be washed out due to degeneracies in pose estimation (Supplementary Fig. S9). For the simulated dataset, we aimed to restrict heterogeneity to a single dimension by selecting conformations along PC1 and limiting variance to a narrow band in PC2. However, because higher-order modes were not filtered, residual variability from these modes in addition to high-frequency motions like side chain dynamics remain present in the data (Supplementary Movie 2). In the future, while multi-dimensional state spaces are realistic in molecular systems, a more careful limitation of the motion in question to a single degree of freedom may be warranted. Additionally, these analyses provided greater insight into the choice of the number of volumes. For example, 80 volumes were initially selected under the assumption that this would increase the likelihood of participants identifying a low-lying state of interest. However, subsequent studies evaluating the number of ground-truth average volumes required to achieve robust distributions under the EMD metric revealed that 80, 40, 20, 16, 10, and even 8 ground-truth–derived volumes all yielded similarly suitable matches (Supplementary Fig. S10). Finally, we note that requesting submissions in the form of volume series rather than atomic models introduced certain challenges, as this format requires standardization steps such as filtering, sharpening, and alignment—procedures that are non-trivial using volumes but essential to ensure fair comparisons. Nevertheless, volumes remain the most widely accessible format for most heterogeneity-focused methods and o”er valuable insights into conformational variability, even when only low-resolution data is available.

In summary, with 41 submissions across both datasets and the development of metrics to assess these submissions, this first conformational heterogeneity challenge in cryo-EM has been informative on many fronts. First, while benchmark datasets based on simulations are immensely valuable [19, 34], this challenge represents the first coordinated e”ort by the community to systematically compare cryo-EM heterogeneity analysis methods on both experimental and simulated datasets. Participants were given the opportunity to optimize their approaches as they saw fit, enabling a realistic assessment of each method’s capabilities. Second, we established a foundation for cryo-EM heterogeneity metrics tailored to specific tasks—capturing motions and distributions—both in the presence and absence of ground truth data. We have brought to the forefront the importance of accurately estimating populations [35], crucial for understanding the thermodynamics of biomolecules. Third, we hope this challenge serves as only the first of many that can drive the field forward towards more confident estimates of conformational ensembles from cryo-EM data. Finally, we are grateful to a community of scientists whose contributions made this project possible and who will remain essential to advancing the field for the many practitioners seeking to confidently use computational methods to extract the information inherent in their cryo-EM datasets.

## Data Availability

All submitted volumes and relative populations, anonymized and pre-processed, are available at https://osf.io/8h6fz/. All analysis code is available at https://github.com/flatironinstitute/Cryo-EM-Heterogeneity-Challenge-1/. The two challenge datasets remain available at https://www.simonsfoundation.org/flatiron/center-for-computational-biology/structural-and-molecular-biophysics-collaboration/heterogeneity-in-cryo-electron-microscopy/.

## Author contributions

YL, YZ, JY, FY, QG, RV, SJ, LD, SJL, HB, SR, MM, VP, JS, ERCC, PS, MAG, AS, DH, JMC, COSS, JRF, and EDZ submitted to the challenge as participants. MAA, DSS, WZ, NG, PC, and SMH devised the challenge, generated the simulated data, and launched the challenge. MK collected and analyzed the electron microscopy data. MAA, GW, DSS, and DH implemented metrics, plotted figures, and developed automated workflow for data processing. MAA, GW, DSS, PC, and SMH devised methodology, coordinated correspondence with co-authors, interpreted results, and jointly wrote the manuscript with feedback from all authors.

## Acknowledgments

The first authors and corresponding authors would like to thank the general support of the Structural and Molecular Biophysics team and the Center for Computational Biology throughout this process. In general we thank our admin team at the Flatiron Institute, including but not limited to Rebecca Sesny who helped with website creation and Christina Julien who helped in compiling co-author information. We are grateful to Reza Paraan and Will Conway for their involvement in discussions at early stages of the project. GW acknowledges various colleagues for helpful discussions: Frederik Kunstner on multiplicative gradient descent for a KL objective; Bruce Shepherd on the linear EMD objective as a min flow problems; Daniel Lee on using Frank Wolfe to solve the Gromov-Wasserstein distance; Minhuan Li on the self-continuity; and Luke Evans on connections to his work on ensemble-reweighting. KDD and GW acknowledge the support of the Government of Canada’s New Frontiers in Research Fund / fonds Nouvelles frontières en recherche du gouvernement du Canada (NFRF), NFRFE-2019-00486. JS acknowledges support from the Medical Research Council (MRC), as part of United Kingdom Research and Innovation (UKRI) (MC UP A025 1013 to Sjors Scheres). AS and MAG are supported in part by the Simons Foundation Math+X Investigator Award and NIH/NIGMS R01GM136780-01. PS acknowledges support from the U.S. National Science Foundation under Award No. STC1231306. The Flatiron Institute is a division of the Simons Foundation. Technical support was provided by the Scientific Computing Core of the Flatiron Institute.

