## Supplement for "The Inaugural Flatiron Institute Cryo-EM Conformational Heterogeneity Challenge"

### Methods

#### 2 Experimental Dataset: Cryo-EM data acquisition, image processing, and refinement

##### 2.1 Sample Preparation and collection of experimental images

The details of the sample preparation protocols for the experimental dataset in this study can be found in ref. [36]. Briefly, 10 mg of thyroglobulin powder (Sigma Aldrich catalog number T9145) was dissolved in a 600 $\mu$L buffer consisting of 10 mM HEPES pH 7.4 and 100 mM NaCl and then purified using gel filtration. The sample was concentrated to 7 mg/mL using centrifugation with a 100 kDa cutoff filter (Millipore). 0.1%(w/v) CHAPS was added to the purified concentrated solution to reduce interactions with the air-water interface. The sample was applied to glow discharged UltrAuFoil R0.6/1 holey gold grids (Quantafoil). Freezing was performed with a Vitrobot (FEI) at 4 °C with 100% humidity, using a blotting time of 9 seconds, a blotting force of 3, a waiting time of 30 seconds and a drain time of 0 seconds. Vitrified grids were imaged on a Krios microscope with a Gatan K2 detector at a magnification of 22,500×, giving a calibrated pixel size of 1.073 Å and a nominal defocus range between -1 and -2 $\mu$m. 1698 movies were collected with a dose of 71.8 $e^-/\text{Å}^2$ in 50 frames over 10 seconds.

##### 2.2 Experimental Data Image Processing

The movies were gain-corrected and subject to motion correction with MotionCorr2 [37]. Patch-CTF with default settings was used to determine the defocus of the micrographs. Particles were picked with a blob picker, and 2D classes were generated from these initial picks. These classes were used as templates for more refined picking. Particles were extracted with a box size of 448 pixels and junk classes were screened out with two rounds of 2D classification and then *ab initio* reconstruction with three classes was performed. Only particles belonging to the highest quality class were selected for further processing. This was followed by a round of homogeneous refinement and another round of 2D classification to further screen out junk particles and particles of a lower quality. Since we intended this challenge to focus on conformational heterogeneity only the highest quality particles from 2D classification or *ab initio* reconstruction were selected for inclusion in further processing. This ensured that as little compositional heterogeneity was present in the dataset as possible.

This careful screening resulted in 33,742 particles. A reconstruction of these particles was produced with homogeneous refinement with an initial low pass filter resolution of 30 Å, a batchsize epsilon of 0.001, an inner window radius of 0.85, an outer window radius of 0.99 and a symmetric noise model. Higher order CTF correction such as beam tilt and trefoil were fit during this reconstruction procedure. This resulted in a 3.3 Å reconstruction. The resulting pose and defocus parameters from this homogeneous reconstruction were used to generate the simulated dataset.

### 3 Simulated Dataset: Conformational distribution and simulated image generation

#### 3.1 Generating atomic structures

##### System building

The starting structure for our molecular dynamics simulation was from PDB ID 6SCJ, which is based on a 3.6 Å resolution cryo-EM volume. The glycans in the PDB structure were removed prior to any modeling. When filling in missing amino acids in the structure, it became apparent that amino acids 848 to 862 were labeled incorrectly as 855 to 869. This was corrected and all missing amino acids were built using Modeler 9.23 [38]. The resulting structure was minimized in implicit solvent by steepest descent to relax the structure at atomic resolution. The system topology was built using Martini 3.0 [39] with secondary structures calculated according to the DSSP program [40]. This coarse-grained structure was then solvated with Martini water and neutralized with 150 mM of sodium chloride.

##### MD simulation protocol and structure processing

Using GROMACS 2023.1 [41], the solvated molecular system was subjected to steepest descent minimization followed by equilibration with 20 femtosecond timesteps for 200 picoseconds. Coarse-grained molecular dynamics runs were carried out in 10 replicates for 6 microseconds each at 350 Kelvin. Coarse-grained conformations were extracted from these simulations in 0.1 nanosecond intervals, resulting in 60,000 structures. These were mapped to atomic resolution using the Backward program [42] followed by minimization in an implicit solvent with NAMD 2.14 [43].

##### Performing PCA on atomic coordinates

Principal Component Analysis (PCA) is often used on molecular dynamics trajectories to determine the largest motions a molecule undergoes during a simulation [44]. To perform this analysis on the backmapped, minimised atomic structures of thyroglobulin, we aligned every structure by minimsing their RMSD to the reference model (PDB: 6SCJ). This removed global translations and rotations from the models. We then defined a data matrix $D$ according the atomic coordinates from the MD trajectory with the $i$th row in the matrix being the flattened cartesian coordinates in the $i$th frame. For $N$ atoms with cartesian coordinates, this becomes

$$D_i = [x_i^1, y_i^1, z_i^1, ..., x_i^N, y_i^N, z_i^N]$$

Hence a trajectory with $N$ atoms and $d$ frames becomes a data matrix $D \in \mathbb{R}^{d \times 3N}$. Using the Singular Value Decomposition (SVD), we can calculate the eigenvectors and eigenvalues of $D$

$$D = USV^T \ , \tag{1}$$

where $U \in \mathbb{R}^{d\times d}$, $S \in \mathbb{R}^{d\times 3N}$, and $\mathcal{V}^T \in \mathbb{R}^{3N\times 3N}$. After SVD, the columns of $\mathcal{V}^T$ will be the principal components of the simulation trajectory. This approach is analogous to the procedure we employ on voxelized volumes for analysis of participants submissions in 6.1.2. Note that during our calculations we have excluded the disordered N and C termini of thyroglobulin (methionine 1 to proline 34 and serine 2723 to lysine 2768) as disordered regions tend to dominate PCA if included due to their high variance [45]. We performed rejection sampling on the backmapped and minimized atomic structures by restricting the range of PC2 according to Fig. 4A.

##### Reweighting the molecular dynamics trajectories

To create a non-trivial but tractable distribution of structures for participants to study, we sampled the frames from the MD trajectories with the following procedure. We first limited the number of unique structures in the simulated dataset, the variance along PC2 was restricted to between -100 and 100 Å and then samples were taken uniformly along PC1, which we chose to treat as the "ground truth heterogeneity coordinate", $z$. This resulted in a selection of 3,861 unique structures in the simulated dataset. Rejection sampling was then used to draw 33,742 samples from these unique structures such that the distribution of structures along PC1 resembled a probability density function defined by the sum of three Gaussian functions (Fig. 4B):

$$\mathrm{PDF}(z) = \frac{1}{Z}\left(\exp\left[-\frac{(z-150)^2}{1500}\right] + \frac{1}{2}\exp\left[-\frac{z^2}{1000}\right] + \exp\left[-\frac{(z+150)^2}{1500}\right]\right), \quad (2)$$

where $z$ represents the coordinate along PC1 and $Z = 5\sqrt{10\pi}(2\sqrt{6}+1) \approx 162.32$ is a normalization constant. This function was generated to reflect realistic properties of biophysical energy landscapes. For example, it exhibits barriers of 4.1 $kT$ between modes, which is thought to be the upper limit for what is preserved during the vitrification of cryo-EM samples [46]. Similarly, the difference in the height of the different Gaussians was chosen to be small at 0.7 $kT$.

#### 3.2 Simulated image stack and volume generation

The realistic *cis*TEM simulator was used to generate the simulated dataset [27]. We used the values of defocus and the rotations from the experimental dataset as inputs to the *cis*TEM simulator. Hence, the ground truth poses and defocus values for the simulated dataset were in fact the values we distributed for the the first dataset, which were obtained using *ab initio* reconstruction and refinement. Meanwhile, the in-plane displacements (translations) for the simulated dataset were sampled on a uniform distribution in X and Y to be between -8.5 and 8.5 Å. Note that the distributed poses for the simulated dataset were obtained by applying the *ab initio* reconstruction and refinement outlined in section 2.2 to the simulated image stack. Exact inputs to the *cis*TEM simulator are outlined in Table 5. The ground truth volumes used were also generated from the 3861 atomic models with the *cis*TEM simulator using the same inputs. Both the experimental and simulated datasets contained 33,742 images.

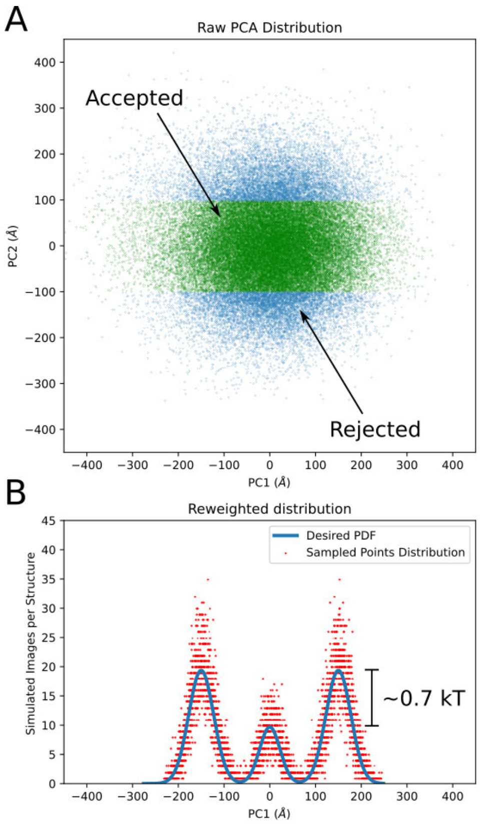


**Fig. 4**: A.) The distribution of all MD frames projected along principal components 1 and 2. Structures with a PC2 greater than 100 Å are excluded from the simulated dataset. From this band, 3861 unique structures are uniformly selected along PC1 for inclusion in the simulated dataset. B.) We draw from these unique structures 33,742 times (the size of the image stack) with rejection sampling to produce a distribution along PC1 that resembles three distinct Gaussians.

Additionally, a second simulated dataset with 20 times the number of images as the other two distributed datasets was also created (for a total of 674,840 images). This larger simulated dataset used the same distribution of molecular structures, poses and defocus values as the smaller simulated stack of images. The only submissions which made use of the larger dataset were Chocolate Chip 2, and Rocky Road 2 and 3. These are denoted with the symbol (*) throughout this article.

#### Simulated image stack validation with *ab initio* reconstruction

*Ab initio* reconstruction with C1 symmetry was carried out in cryoSPARC to a target resolution of 6 Å, with 200 and 300 iterations before and after annealing, respectively. Heterogeneous refinement was performed by using cryoSPARC 4.1 defaults with C2 symmetry. This was followed by local refinement with C2 symmetry with cryoSPARC defaults. The set of defoci and poses resulting from the obtained reconstruction process is what we distributed to participants as reasonable starting values for the simulated dataset.

##### 3.3 Ground Truth Controls

We defined different types of controls to represent the ground truth. We ordered the 3,861 volumes by ground truth heterogeneity coordinate (defined in section 3.1), then divided the space into 80 disjoint sets of equal interval. For *Averaged GT*, in each subset we took the Euclidean mean of the volumes and population weights to form 80 smoothed volumes and 80 smoothed weights. For *Sampled GT*, the structure from each interval with the highest population was taken. This resulted in 80 volumes which directly represent 1224 particles in the simulated image stack.

##### 3.4 Ordering of volumes

On the challenge website we read: *Please use your method of choice to analyze these data and provide results in the form of 80 volumes (at* 2.146 *Å/voxel aligned to the reference volume) with the relative population of each, and their coordinates along the dominant degree of freedom (finely sampling the direction of greatest conformational change).* However, most participants ordered their volumes without providing separate information about their spacing. A few groups submitted volumes in a seemingly random order (cf. submitted population in Fig. 3B). Of note, none of our metrics required the use of the submitted order of the volumes, with the minor exception of visualization tasks in the following cases: the middle volume (i.e. 40th) is used to visualize results in section 1.4, and for the EMD/KL metric the five nearest neighbor window is used to smooth the plotting (section 6.5.3).

#### 4 Heterogeneity inference methods used by participants

Below, participants have provided the details of the methods used for generating the 80 volumes and their respective populations for each submission. To maintain anonymity, we do not associate them with their corresponding ice cream flavor names. Participants were informed of their ice cream flavor in individual discussions with the analysis team.

##### 4.1 Method A

The method used is a subspace method. A PCA-based algorithm was employed to embed the dataset, and a kernel regression estimator was applied to generate volumes. The reported populations were computed using a deconvolved kernel density estimator derived from the embedded dataset along the first principal component. More details on the method are available at [29]. Poses were not re-estimated.

#### 4.2 Method B

The given poses were used for training the DynaMight VAEs. The particles with the corresponding poses were then used for training following [47]. Here the poses were refined *via* gradient descent within the network training. After training the latent space was computed, the dimensionality reduced by PCA. A histogram with 80 bins along the first component was computed to extract the populations and the associated volumes.

#### 4.3 Method C

Particles were first aligned to estimate their relative poses with respect to a single consensus model. These aligned particles, together with the consensus model, were then used to train a newer more efficient version of the neural network originally introduced by Chen and Ludtke [48] (e2gmm.py). Particles with known orientations were represented as 3D Gaussians, where shifts in Gaussian amplitudes and positions reflected local structural variations of the protein. The trained neural network embedded these Gaussian representations into low-dimensional latent spaces—four dimensions for dataset 1 and one dimension for dataset 2. To generate volumetric reconstructions with consistent resolution, 80 cluster centers were selected in the latent space: evenly spaced along a curved trajectory identified in the 2D PCA projection of the 4D latent space for dataset 1, and along the single latent dimension for dataset 2. From each cluster center, a subset was formed by including the 3,000 nearest neighboring particles in the latent space. The population distribution of all subsets was then computed based on the normalized local density at their corresponding cluster centers.

#### 4.4 Method D

A linear subspace model of cryo-EM density was fit to particle data, with each particle being represented as a consensus density plus a weighted sum of components [49]. The estimated component weights provide a latent embedding of the particles, from which populations were separated by binning along the direction of maximum variance.

#### 4.5 Method E

A neural network model representing 3D continuous deformation of a canonical volume was fit to the particle data [50]. The weights of the deformation network, the density values of the canonical volume, and the latent embedding vectors of each particle were jointly optimized. After training the model, the latent embedding of each particle (which encodes a specific 3D deformation field for each particle) was transformed into a secondary latent embedding space using another neural network. This network was trained so that distances in the secondary latent space would correspond to distances (i.e. Euclidean norm) in 3D deformation space. Particle latent positions in the secondary space were subject to PCA and then binned along the first principle component to produce populations.

#### 4.6 Method F

We employed the ManifoldEM approach [51, 52]. In short, ManifoldEM involves four sequential data processing steps: (1) pre-processing and binning of the particles into similar projection directions; (2) embedding of the binned particles by Diffusion Map for each projection direction; (3) rendering of 2D Non-linear Laplacian Spectral Analysis (NLSA) movie frames along the conformational coordinates; (4) patching of all movie frames consistently across all projection directions. To obtain the 3D movie of conformations, we reconstructed 3D volumes with relion_reconstruct. The poses were provided by the organizers of the Heterogeneity Challenge. The relative populations were determined from the ten most populated projection directions.

#### 4.7 Method G

Particles were initially refined, providing pose and in-plane shifts using an angular consensus. The refined particles were then used to train a neural network to predict a set of coefficients to be used in conjunction with the Zernike3D basis described in [53]. The network was trained with a learning rate of $1 \times 10^{-4}$ for each dataset, batch sizes of 8 and 64 images for the first and second datasets, respectively, and pose and CTF decoupling architectures. After training, the coefficient space was recovered and subjected to a PCA analysis. We then divided the first principal component into 80 groups with the same bin length to count the number of coefficients falling into each group, thereby recovering the population associated with each state. Additionally, we took the middle point of each bin to recover a new set of coefficients, considering only the first principal component, which was then used to recover the deformation field needed to bend the original reference volume towards the conformation associated with a given population.

#### 4.8 Method H

Particles were initially refined, providing pose and in-plane shifts using an angular consensus. The refined particles were then used to train a neural network to predict a set of coefficients to be used in conjunction with the Zernike3D basis described in [53]. The network was trained with a learning rate of $1 \times 10^{-4}$ for the second dataset, a batch size of 64 images, respectively, and pose and CTF decoupling architectures. After training, the coefficient space was applied in ZART [54] as a non-linear alignment to correct for the motion artifacts present in the original reference volume, thus improving its local resolution. The corrected volume was then used to compute a new set of coefficients, which were used again to improve the quality of the corrected volume further. This iterative process was repeated until convergence (which took around five iterations), yielding a final corrected volume with an improved coefficient space. The improved coefficients space was further analyzed to extract the populations and the associated volumes. The analysis involved dimensionality reduction of the spaces using UMAP [55], followed by principal curve analysis to determine the optimal curve that traverses the landscape. Once the principal curve was determined, the landscape was projected on it, and the curve was split into 80 segments. By counting the number of projected points in each segment, it was possible to extract the relative populations and their associated volumes, which were obtained by decoding the midpoint of each segment with the trained networks.

#### 4.9 Method I

HetSIREN analysis [56] started by refining the particles' provided pose and in-plane shifts using an angular consensus. The refined particles were then used to train HetSIREN neural network with a learning rate of $1 \times 10^{-5}$ for each dataset, batch sizes of 8 and 64 images for the first and second datasets, respectively, and pose and CTF decoupling architectures. After training, the conformational latent space was predicted and further analyzed to extract the populations and the associated volumes. The analysis involved dimensionality reduction of the spaces using UMAP [55], followed by principal curve analysis to determine the best possible curve that traverses the landscape. Once the principal curve was determined, the landscape was projected on it, and the curve was split into 80 segments. By counting the number of projected points in each segment, it was possible to extract the relative populations and their associated volumes, which were obtained by decoding the midpoint of each segment with the trained network.

#### 4.10 Method J

A cryoDRGN model [57] was trained on both datasets of refined particles using an encoder-decoder architecture of 1024×3 with a latent variable dimension of 8 for 50 epochs. Population counts were estimated by histogramming the latent embeddings along the first principal component axis of the latent space with 80 equally spaced bins, excluding outliers in the 1 and 99 percentiles. Representative volumes were generated from the center of each bin using the trained decoder network.

#### 4.11 Method K

Starting from the particle poses and the model provided by the challenge organizers, this method used data-driven coarse-grained (C-alpha Go) MD simulations accelerated with normal modes (MDSPACE [58]), resulting in one coarse-grained conformational model per particle image. The method allowed the model to change its pose during each MD simulation to adapt to the pose of the particle in the image, which is closely related to the particle pose refinement [58]. Two MDSPACE iterations were performed, each with 40 picosecond C-alpha Go MD simulations and force constant of 6000 kcal/mol. In the first iteration, MD simulations were combined with 13 normal modes. In the second iteration, normal modes were replaced by 6 principal component vectors obtained by PCA at the end of the first iteration. Such MDSPACE iterative strategies were shown to help refining the conformations [58]. The coarse-grained models obtained with MDSPACE (one model per particle) were rigid-body aligned before performing PCA or UMAP analysis to embed them in a common low-dimensional space [58]. Currently, the software is not optimized for extracting populations, but it provides several options for interactive and automatic splitting of the PCA/UMAP space into local regions to calculate the average coarse-grained models, density (volumes) from these models, and 3D reconstructions from images in these regions [59]. Here, the partitioning was done automatically by splitting the first PCA/UMAP axis into 80 segments. The submitted populations were obtained by counting the number of points (the associated models or images) in each segment. A relatively small number of images per segment (large number of segments) resulted in very noisy 3D reconstructions. Therefore, we submitted the average coarse-grained models converted into volumes.

##### 4.12 Method L

CryoSTAR integrates atomic-model priors with single particle images with given poses (provided by other software) by first learning a low-dimensional latent embedding under structural regularization to sample coarse-grained conformers. An MLP then decodes these latents into volumetric density maps, and conformer populations are quantified by projecting latent vectors onto principal components followed by clustering [60].

#### 5 Submission pre-processing

Before running any of the comparison pipelines, we preprocessed the submissions so that all volumes had the same pixel size, box size, and alignment. For the pixel and box sizes, we padded or cropped volumes, and if needed volumes were downsampled so that they had a box size of 224, and a voxel size of 2.146Å. Only volumes with a smaller voxel size were downsampled. Downsampling was performed with the Hartley transform, following the implementation in cryoDRGN [57]. Some submitted volumes had the wrong handedness, in this cases we flipped their handedness before aligning. The volume alignment is described below.

##### 5.1 Volume alignment

The alignment of the submitted volumes is the most delicate step in the preprocessing pipeline, as differences in alignment can easily perturb the results in our metrics. Before aligning a submission, we remove the dust in each volume by setting to zero all pixels with value lower than the $p$-th percentile, were $p$ was defined on a per-submission basis. Then, for each submission, we compute the average volume, apply an initial manual alignment defined by visual inspection, and then run the BOTAlign algorithm [61] to refine the alignment as implemented in Aspire [62]. The obtained rotation matrix is then applied to all volumes in the given submission. The manual alignment is performed to avoid the BOTAlign algorithm getting stuck in local optima, and reduce its computational cost. The BOTAlign algorithm was run using the WEMD loss, and the initial phase was run with the downsampling box size was set to 64 for a maximum of 400 iterations. The refinement phase was run with the default parameters defined in Aspire [62]. The submissions were aligned to 1) the consensus reconstruction for the experimental dataset, and 2) the average of the ground truth volumes for the simulated dataset. We chose not to do an all-to-all alignments (3861 x 80 = 308,880) so as to preserve the relative alignments in the submissions. All volume alignments were visually inspected in Chimera at the end. The transformations performed on each submission are recorded in Supplementary Tables 1, 2, 3, and 4.

#### 5.2 Masking

A loose mask was generated to remove the background of user-submitted volumes (Fig. 5), and was used when computing volume-to-volume distances. Values were completly excluded from outside the mask (i.e. not thresholded to zero), and the computation only used voxels inside the mask. Note that during each Zernike3D volume-to-volume distance computation there is a bespoke mask generated, which is explained in section 6.2.11.

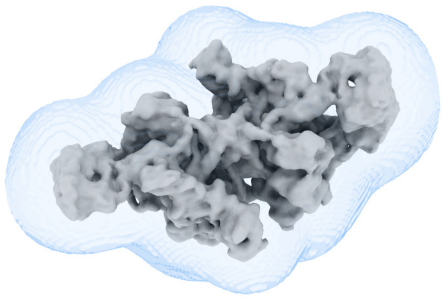

**Fig. 5**: The consensus refinement from the simulated dataset pictured within the dilated mask.

### 6 Metrics

#### 6.1 Principal Component Analysis

In this section, we describe the PCA-based approach for comparing the submissions. Using PCA allows us to compare one volume series to another volume series and evaluate their similarity both quantitatively and qualitatively. We begin by outlining the preparation process for submissions prior to PCA analysis. After the preparations, we compute the PCA decompositions of the submissions *via* randomized SVD [63]. Finally, we use these decompositions to find a common embedding, which allows us to visualize all volume series in a low-dimensional space (see Algorithm 1).

##### 6.1.1 Preparing submissions for PCA

We start with the submissions processed as described in Section 5. Next, we apply a dilated mask to all volumes, remove the dust in the volumes by setting to zero all voxels with intensity value below the 97th percentile, and downsample to a box size of 128. Lastly, the $\ell_2$-norm of all downsampled volumes is set to 1. Additionally, we compute the radial power spectrum for each submission and use it to define a whitening filter. To bring all submissions to the same resolution, we compute a common envelope by comparing the power spectra of all submissions as follows. Let $P_{ik}$ be the power spectrum of the $i$-th submission at the $k$-th frequency shell. The common envelope is defined as

$$\tilde{P}_k = \min_{i \in N_{sub}} P_{ik} \ , \tag{3}$$

where $N_{sub}$ is the number of submissions to be compared.

We then proceed by whitening the submissions and applying the common envelope. This approach was chosen as the metric was particularly sensitive to differences in resolution between submissions. For the simulated dataset, we also compare submissions to the ground truth volumes *via* the two ground truth mock submissions. Although these mock submissions are also considered when computing the common power spectrum, they have a relatively high resolution compared to other submissions, and thus do not affect its computation in practice.

##### 6.1.2 Performing PCA on volume series

PCA consists of computing the eigendecomposition of the covariance matrix of a given data matrix $C = (X - \overline{X})^T(X - \overline{X})$, where $X \in \mathbb{R}^{n\times d}$ is the data matrix with mean $\overline{X}$. In the case where the data is highly dimensional $d \gg n$, this decomposition can be computed efficiently through the Singular Value Decomposition (SVD) of the mean-removed data matrix $D = X - \overline{X}$.

In our case, we define the data matrix associated with submission $i$ ($X_i$) as a stack of flattened volumes. That is, the $j$-th row of the data matrix, $[X_i]_j$, is the flattened $j$-th submitted volume. We remove the mean from this data matrix $D_i = X_i - \overline{X_i}$ and compute its SVD

$$D_i = U_i S_i \mathcal{V}_i^T \ , \tag{4}$$

where $U_i \in \mathbb{R}^{n\times n}$, $S_i \in \mathbb{R}^{n\times d}$, and $\mathcal{V}_i \in \mathbb{R}^{d\times d}$. The matrix $S_i$ is diagonal, with diagonal elements satisfying $\sigma_1 \geq \sigma_2 \geq ... \geq 0$. If the rank of the matrix is $q$, then $\sigma_{q+1} = 0$. This fact is usually used to define the economy-SVD, which simply consists on truncating $U_i$, $S_i$, and $V_i$ such that $U_i \in \mathbb{R}^{n\times q}$, $S_i \in \mathbb{R}^{q\times q}$, and $\mathcal{V}_i \in \mathbb{R}^{d\times q}$.

The eigendecomposition of the covariance matrix can be easily obtained from the SVD as

$$C_i = D_i^T D_i = \mathcal{V}_i S_i^2 \mathcal{V}_i^T, \tag{5}$$

where the right singular vectors $\mathcal{V}_i$ correspond to the eigenvectors of the covariance matrix, and the squared singular values $S_i^2$ to its eigenvalues.

A useful feature of SVD is that the matrix $\mathcal{V}_i$ works as a projector of volumes to a lower-dimensional subspace $\mathbb{R}^d \to \mathbb{R}^q$. In the following subsections, we show how this can be used to embed all submissions into a common (lower-dimensional) subspace. We can also use this to study how other submissions behave in a given latent space. This will be used to compare submissions to the ground truth volumes. In our implementation, we compute the SVD through the randomized algorithm developed in ref. [63] available in PyTorch [64].

##### 6.1.3 Using PCA to compare submissions

To compare submissions, we use the percentage of captured covariance (PCV) introduced in ref. [29]. This metric computes how much variance is captured by a set of eigenvectors, given a reference basis. In our case, we compare submissions to each other. Let the decomposition computed for submission $i$ define the reference basis, with eigenvectors and eigenvalues given by its SVD: $S_i, \mathcal{V}_i$. Similarly, let submission $j$ and its decomposition define the basis that we will test against the reference. Then the PCV of submission $i$ by submission $j$ is given by

$$PCV(j,i) = \frac{\|\mathcal{V}_j^T \mathcal{V}_i S_i\|_F^2}{\|\mathcal{V}_i^T \mathcal{V}_i S_i\|_F^2} = \frac{\sum_{l,k}[S_i]_k^2([\mathcal{V}_j]_l^T[\mathcal{V}_i]_k)^2}{\sum_k [S_i]_k^2} \,, \tag{6}$$

with $\|\cdot\|_F$ denoting the Frobenius norm. The second equality in Equation 6 shows that the metric is simply a weighted sum of all the possible inner products between the reference basis ($\mathcal{V}_j$) and the basis being evaluated ($\mathcal{V}_i$), with the weights being the singular values of the reference basis. This metric has 1 as its maximum value, which means that submission $j$ captures all the variance in submission $i$. This metric only works for truncated SVD's, as a full rank basis would always give 1 due to the $\mathcal{V}$ matrices being unitary.

###### 6.1.4 Using PCA to compute a common embedding between submissions

Now we proceed to describe a heuristic approach for visualizing the different submissions in a reduced-dimensional space. We can achieve this by considering that the eigenvectors $\mathcal{V}_i$ of the covariance matrix serve as a projector to a lower-dimensional space, with dimensions given by $q$, the rank of the mean-removed data matrix. However, comparing the projection of all methods to all the different subspaces is infeasible. For this reason, we use the eigenvectors computed for each submission to compute a new SVD, and thus define a common lower-dimensional subspace.

We start by weighting all eigenvectors by their associated eigenvalue $[S_i]_k[\mathcal{V}_i]_k$ for $k = 1, \ldots, q$ and $i = 1, \ldots, N_{sub}$, where $N_{sub}$ is the number of submissions. Then we stack all the weighted eigenvectors obtained for all submissions to define a new matrix $Y \in \mathbb{R}^{(N_{sub}q)\times d}$. We weight the $\mathcal{V}_i$s to avoid eigenvectors associated with small singular values to take over the computation of the basis. Similar as before, we compute the SVD for $Y$ after removing its mean:

$$Y - \overline{Y} = \tilde{U}\tilde{S}\tilde{\mathcal{V}}, \tag{7}$$

where $\tilde{U}, \tilde{S} \in \mathbb{R}^{(N_{sub}q)\times(N_{sub}q)}$, and $\tilde{\mathcal{V}} \in \mathbb{R}^{(N_{sub}q)\times d}$. Then we can use the projector properties of $\tilde{\mathcal{V}}$ to project all the submissions onto this common subspace.

##### 6.2 Volume-to-volume distances

Unlike atomic models where a correspondence can be found between points, volumes have voxel intensity spread out over spatial neighborhoods, and different internal weighting of intensity depending on each participant's heterogeneity inference method (see section 4), or forward model [27]. In order to quantify how "close" a volume is to another conformationally heterogeneous volume, we experimented with several different types of volume-to-volume distances. Our criteria included computational

**Algorithm 1** Obtain SVD for a submission

**Require:** submission dictionary containing an id, and volumes

normalize ← (threshold, mask, downsample, normalize power spectrum, set $\ell_2$-norm to 1)
volumes ← normalize(submissions["volumes"])
volumes ← volumes - mean(volumes)
$U$, $S$, $\mathcal{V}^*$ ← SVD(volumes)

**return** $U$, $S$, $\mathcal{V}$

efficiently, suitability for quantifying displacement of mass (i.e. a proxy for RMSD of atomic positions), robustness to small perturbations in conformational heterogeneity (largely from high resolution details in the ground truth volumes from side chain fluctuations), and invariance to volume sharpening and point-wise multiplicative and additive scaling of voxel intensity and pose. Specifically we used volume-to-volume distances based on mean squared error and correlation of voxel intensities, the log probability of a 3D white Gaussian noise model with marginalization, Fourier shell correlation averaged across frequencies, optimal transport (Wasserstein, Sliced-Wasserstein, Procrustes-Wasserstein and Gromov-Wasserstein), and a deep learning volume deformation field based on the Zernike polynomials (Zernike3D). Results are shown in section 6.4. Unless stated otherwise, all volumes were masked with a wide dilated mask and normalized before computing the volume-to-volume distance.

##### 6.2.1 Volume normalization

For volume-to-volume distances we normalized volumes using Eq. (8). For optimal transport volume-to-volume distances in sections (6.2.7, 6.2.8, 6.2.9, 6.2.10) we further normalized the volumes by Eq. (9) if there was negative voxel intensity, otherwise by Eq. (10):

$$V^{\text{median-std}} = \frac{V - \text{median}(V)}{\sigma(V)} \tag{8}$$

$$V^{\text{min-prob}} = \frac{V - \min(V)}{\sum_i [V - \min(V)]_i} \tag{9}$$

$$V^{\text{prob}} = \frac{V}{\sum_i [V]_i} \tag{10}$$

$$V^{\text{mean-std}} = \frac{V - \text{mean}(V)}{\sigma(V)} \tag{11}$$

##### 6.2.2 Mean squared error

The mean squared error (MSE) was computed *via* the Frobenius norm, which is equivalent to the squared $\ell_2$ norm of the flattened (turned into 1D vector, denoted by vec[ ]) 3D volumes.

$$d_{\mathrm{mse}}(i,j) = ||V_i - V_j||_F = ||\mathrm{vec}[V_i] - \mathrm{vec}[V_j]||_2 \tag{12}$$

where $i$ and $j$ are for different volumes and $||\cdot||_F$ is the Frobenius norm.

##### 6.2.3 Real-space voxel correlation

The (negative) correlation was computed *via* the inner product between the two volumes.

$$f_{\mathrm{corr}}(i,j) = -\mathrm{vec}[V_i]^T \mathrm{vec}[V_j] \tag{13}$$

Note that pre-normalization with mean-std (Eq (**??**)) makes this metric invariant to global multiplicative and additive scalars $N, \mu \in \mathbb{R}$ (see Eq. 18 in [65]). However we still normalized with Eq (8); when the mean and median values are similar the correlation is approximately invariant.

$$\begin{aligned}
&- \mathrm{vec}[(N_i V_i + \mu_i)^{\text{mean-std}}]^T \mathrm{vec}[(N_j V_j + \mu_j)^{\text{mean-std}}] &(14)\\
&= - \mathrm{vec}[V_i^{\text{mean-std}}]^T \mathrm{vec}[V_j^{\text{mean-std}}] &(15)\\
&\approx - \mathrm{vec}[V_i^{\text{median-std}}]^T \mathrm{vec}[V_j^{\text{median-std}}] &(16)
\end{aligned}$$

##### 6.2.4 BioEM3D: marginalized mean squared error

We extended the marginal likelihood from [66] (Eq. (10) in their supplement), which was originally used to compare two images, to be a distance between two volumes. In brief, under a Gaussian model, the likelihood analytically marginalizes out additive and multiplicative scaling with a flat prior, and then marginalizes out the variance around the maximum using the saddle point approximation. Thus, we define the BioEM3D volume-to-volume distance as,

$$\begin{aligned}
& d_{\mathrm{BioEM3D}}(V_i, V_j) \overset{\mathrm{def}}{=} -\log p_{\mathrm{BioEM3D}}(V_i, V_j) \\
\text{where}\quad & p_{\mathrm{BioEM3D}}(V_i, V_j) = \sqrt{\pi}(2\pi e)^{1-n_{\mathrm{vox}}/2} \\
& \quad \times \left[n_{\mathrm{vox}}(C_{ii}C_{jj} - C_{ji}^2) + 2C_j C_{ji} C_i - C_{ii}C_j^2 - C_{jj}C_i^2\right]^{3/2 - n_{\mathrm{vox}}/2} \\
& \quad \times \left[(n_{\mathrm{vox}} - 2)(n_{\mathrm{vox}} C_{ii} - C_i^2)\right]^{n_{\mathrm{vox}}/2 - 2},
\end{aligned} \tag{17}$$

and where $n_{\mathrm{vox}}$ is the total number of voxels (e.g. $n^3_{\text{box size}}$ with no masking); and $C_{(\cdot)}$, and $C_{(\cdot,\cdot)}$ are the sum and correlation between volumes $V_i$ and $V_j$, following Eqs. (5)-(9) in the supplement of [66].

##### 6.2.5 Fourier shell correlation (FSC) & Resolution assessment

Since we have GT volumes for the simulated dataset, we compute the Fourier shell correlation (FSC) comparing each of the 80 submitted volumes $\{V_j^{(u)}\}_{j=1}^m = 80$ to each the 3861 GT volumes $\{V_i^{(t)}\}_{i=1}^m = 3861$, and estimate the resolution at the 0.5 threshold: $r^{i,j} = \arg\max_r \mathrm{FSC}_r(V_i^{(t)}, V_j^{(u)}) = 0.5$. We then take the best resolution for each submitted volume:

$$r^j_{\text{best}} = \max_i r^{i,j} \ . \tag{18}$$

Therefore, for each submission we have the set $\{r^j_{\text{best}}\}_{j=1}^{n=80}$ (shown in Fig. 2**E**).

##### 6.2.6 Volume-to-volume FSC distance

The Fourier shell correlation is invariant to volume sharpening, as it computes the normalized correlation $\in [-1, 1]$ at some frequency band $[k_-, k_+)$. Any multiplicative b-factor weightings $w_i, w_j$ are canceled out.

$$\mathrm{FSC}_k(V_i^{(t)}, V_j^{(u)}) = \frac{\mathrm{Re}\big[[\hat{V}_i^{(t)}]_k^\dagger [\hat{V}_j^{(u)}]_k\big]}{||[\hat{V}_i^{(t)}]_k||_2 ||[\hat{V}_j^{(u)}]_k||_2} \tag{19}$$

$$= \frac{\mathrm{Re}\big[[w_i\hat{V}_i^{(t)}]_k^\dagger [w_j\hat{V}_j^{(u)}]_k\big]}{||[w_i\hat{V}_i^{(t)}]_k||_2 ||[w_j\hat{V}_j^{(u)}]_k||_2} \tag{20}$$

where $\hat{V}$ is the discrete Fourier transform of $V$, $\dagger$ is the complex transpose, $[V]_k$ is the vector of elements in $V$ at radius $k \in [k_-, k_+)$, and Re is the real-spared projection operator, and $||\cdot||_2$ is the $\ell_2$ norm.

A distance based on FSC was defined for a spectral band $S_k = \{k_{\text{min}}, ..., k_{\text{max}}\}$ as

$$d_{\text{FSC}}(V_i^{(t)}, V_j^{(u)}) = 1 - \frac{1}{|S_k|} \sum_{k \in S_k} \mathrm{FSC}_k(V_i^{(t)}, V_j^{(u)}) \tag{21}$$

which essentially computes the "area under the FSC curve". The maximum of this distance is 1, when the $\mathrm{FSC}_k$ is 1 over all frequencies. Note that in our analysis we observed it to be always greater than zero, but its theoretical minimum is $-1$ when $t_i = -u_j$.

##### 6.2.7 Wasserstein

We compared volumes via the Wasserstein distance, with a squared Euclidean cost matrix, and the marginals being the top-$k$ grid intensities with the most mass ($k$ typically in the high hundreds to low thousands) after down-sampling (typically to a box size in the range 32-64) via Fourier cropping. We used the unregularized Earth Mover's Distance algorithm implemented in Python OT [67] to determine the transport plan, which is $O(k^3)$. We took $k$ high enough to qualitatively visually distinguish the conformational motion across the trajectory. Excessive down-sampling makes the states difficult to distinguish. For a fixed $k$, insufficient down-sampling results in excessive mass being excluded, and concentrates on high intensity regions that do not spatially displace. Each marginal was normalized by Eq. (8) followed by (10). For the results in Fig S4 each volume was down-sampled to a box size of 32 and the top-500 intensity voxels were used. Thus we used the squared Euclidean distance for the cost matrix $(500 \times 500)$.

##### 6.2.8 Sliced-Wasserstein

We computed the Sliced-Wasserstein (SW) to approximate the Wasserstein distance. Volumes were assumed to be aligned, and random projections were generated (same projection shared by volume pairs) from a uniform distribution over rotations. Volumes were interpolated on a rotated regular grid with `torch.grid_sample` and then projected to a regularly spaced 1D array by summing along two axes. The SW was approximated by computing the $\ell_2$-norm of the cumulative distribution function (cdf) of projections. Note that we did not invert the cdf, since we found that this cheap approximation suffices for our purposes; see Remark 2.30 in [68] and Chapter 2 of [69]. We benchmarked convergence as a function of initial volume down-sampling and the number of projections. At a box size of 64, increasing the number of projections from several hundred to a several thousand projections did not show noticeable improvement, and the standard deviation was around two orders of magnitude below the mean. For the results in Fig S4 we used 300 random 1D projections of the volume down-sampled to a box size of 64.

##### 6.2.9 Procrustes-Wasserstein

The Wasserstein distance, as explained in section 6.2.7 is not invariant to the local alignment of the two volumes, and we therefore iteratively optimized the alignment via an SVD on the correlation of the location of the points through the transport plan. This iterative algorithm is known as the Procrustes-Wasserstein [70, 71]. For the results in Fig S4 we treated the volumes in the same way as in section 6.2.7.

##### 6.2.10 Gromov-Wasserstein

We used another optimal transport distance that is invariant to rotations (and reflections) called Gromov-Wasserstein (GW)

$$d_{\mathrm{GW}}(a,b) = \min_{\Gamma \in \Pi} \sum_{ijkl} |C^{(a)}_{ij} - C^{(b)}_{kl}|^2 \Gamma_{ik} \Gamma_{jl} \tag{22}$$

where $C^{(a)}$ is some cost in the space of volume $a$, and $C^{(b)}$ is some cost in the space of volume $b$, and $\Gamma$ is a transport plan of voxel intensities from volumes $a$ and $b$. We used the Euclidean distance for $C_{(a)}, C_{(b)}$, which is invariant for arbitrary rotation and shift (desirable); and overly invariant to mirroring (undesirable). As with the Procrustes-Wasserstein, the transport plan $\Gamma$ belong to the convex set $\Pi$ of non-negative matrices with fixed marginals that match the voxel intensities. As noted in Figure S9, the heterogeneity and pose are not identifiable, due to the (pseudo)-C2 symmetry of thyroglobulin, and the direction of the motion of heterogeneity. Most submissions could be divided into half, with one half set a 180 degree rotation of the other half set. Intuitively, GW computes an all-by-all self distance, and then matches points such that the sum of the squared difference of the self-edges is minimal. For symmetric objects the transport plan can match nodes between the asymmetric units. However, in our case, we only use the value of GW, and do not need the transport plan to match the corresponding asymmetric unit in the other volume.

We computed the GW distance using the local solver `ot.gromov_wasserstein` in Python OT [67], which implements a conjugate gradient solver. To normalize the marginals we used the same down-sampling, top-k, and normalization approach as in sections 6.2.7 and 6.2.9. To overcome what we think may be numerical instabilities we exponentiated the (squared Euclidean) cost matrix by 4.

Secondly, we extended the Gromov-Wasserstein objective in [72] to apply to weighted point sets (same number of points). Previously the optimization was over unweighted points and leveraged the fact that the doubly stochastic matrices is a convex set. In our case, the set $\Pi$ of joint probabilities with fixed marginals is also convex, although the vertices are no longer the permutation matrices in general [73]. As noted in [72], when the cost matrices $C^{(\cdot)}$ are the squared Euclidean distance, solving the GW problem is equivalent to solving the quadratic program

$$\begin{aligned} &\min_{\Gamma\in\Pi} -\|2X\Gamma Y^T\|_F^2 - \langle L, \Gamma\rangle + c_0 \\ X &= (x_1, \dots, x_n) \in \mathbb{R}^{3\times n}, \quad \text{and likewise for } Y \\ \text{with } L &= 2(\mu_y^T \mathbf{1}_y) m_x m_y^T - 4 m_x \mu_y Y^T Y - 4 X^T X \mu_x m_y \\ c_0 &= \frac{\langle C_x, C_x\rangle + \langle C_y, C_y\rangle - \mathbf{1}^T m_y \mathbf{1}^T m_x}{2} \\ m_x &= (||x_1||^2, \dots, ||x_n||^2)^T, \quad \text{and likewise for } m_y \end{aligned} \tag{23}$$

We solved Eq. (23) by Frank-Wolfe (another local optimization algorithm). Instead of projecting onto the joint distribution with fixed marginals (as in projected gradient descent), Frank-Wolfe performs a linear assignment, and in our case we utilize the Earth Mover Distance solver `ot.emd`. The gradient term for the linear assignment requires two matrix multiplications that are $O(n^2 d + nd^2)$, where $d = 3$ and $n$ is the number of points. Furthermore, the optimal step size $\eta$ can be found analytically at each step, and the trace efficiently computed with a sum of the Hadamard product. We clip the step size to be between 0 and 1, so that the update keeps $\Gamma$ in its convex domain $\Pi$.

###### 6.2.11 Zernike3D

As an alternative to similarity measurements that rely on comparing voxel values between a pair of volumes, we developed a distance that measures how much we should warp a given volume to match the structural features present in another. This

**Algorithm 2** Optimization of $\min_{\Gamma\in\Pi} -\|2X\Gamma Y^T\|_F^2 - \langle L, \Gamma\rangle + c_0$ via Frank-Wolfe.

**Input:** Point locations $X, Y \in \mathbb{R}^{3\times n}$; marginals $\mu_x, \mu_y \in \Delta^{n-1}$; Point norms $m_x, m_y \in \mathbb{R}^n_{\geq 0}$

**Initialize:** $\Gamma_0 \in \Pi$, where $\quad \Pi = \{\Gamma : \Gamma\mathbf{1}_x = \mu_y,\ \mathbf{1}_y^T\Gamma = \mu_x^T\}$

Compute:

$$L = 2(\mu_y^T\mathbf{1}_y)m_x m_y^T - 4m_x\mu_y Y^T Y - 4X^T X\mu_x m_y$$

**for** $t = 0, 1, 2, \ldots$ until convergence **do**

  Compute gradient:

$$\nabla_{\Gamma_t} f(\Gamma_t) = -8X^T X\Gamma_t Y^T Y - L$$

  Solve linear assignment:

$$S_t = \arg\min_{S\in\Pi}\langle \Gamma_t, \nabla_\Gamma f(\Gamma_t)\rangle$$

  Compute step size:

$$\eta_t = \frac{-8\,\mathrm{tr}[(S_t-\Gamma_t)^T X^T X\Gamma_t Y^T Y] - \mathrm{tr}[L^T(S_t-\Gamma_t)]}{8\,\mathrm{tr}[(S_t-\Gamma_t)^T X^T X(S_t-\Gamma_t)Y^T Y]}$$

  Update:

$$\Gamma_{t+1} = (1-\eta_t)\Gamma_t + \eta_t S_t$$

**end for**

metric quantifies the mass displacement from a reference state to a target volume and condenses such displacement into a similarity metric for a given volume pair.

The displacement can be expressed as a deformation field, where each component determines where to move a given voxel in space. To compute this field efficiently, we use the Zernike3D basis [74], which has already been applied to compute similarity among different volumes following similar principles. Given a volume pair $V_i$ and $V_j$, finding the deformation field that aligns $V_2$ towards $V_1$ reduces to minimizing:

$$\|V_i(\mathbf{r}) - V_j(\mathbf{r} + \mathbf{g}(\mathbf{r}))\|^2, \tag{24}$$

where $\mathbf{r} \in \mathbb{R}^3$ denotes the voxel position and $\mathbf{g}(\mathbf{r})$ is the deformation field at that location.

The Zernike3D basis enables the approximation of an arbitrarily complex deformation field from a reduced set of coefficients

$$\mathbf{g}(\mathbf{r}) = \sum_{l=0}\sum_{n=0}\sum_{m=-l}^{l} \lambda_{l,n,m} Z_{l,n,m}(\mathbf{r}), \tag{25}$$

where $\lambda_{l,n,m}$ are the coefficients indicating the importance of each basis function, and $Z_{l,n,m}(\mathbf{r})$ are the Zernike3D basis functions composed of Zernike polynomials and spherical harmonics, as described in the original work.

Once the coefficients $\lambda_{l,n,m}$ have been optimized, the full deformation field can be reconstructed and used to compute a RMSD from its vector field. This RMSD serves as the distance between the two volumes (with a distance of zero for a deformation of all zeros everywhere), and is given by:

$$\text{Zernike3D} \stackrel{\text{def}}{=} \sqrt{\frac{1}{n_{\text{pix}}^3} \sum_{i=1}^{n_{\text{pix}}^3} g_x^2(\mathbf{r}_i) + g_y^2(\mathbf{r}_i) + g_z^2(\mathbf{r}_i)} \tag{26}$$

where $\{\mathbf{r}_i\}_{i=1}^{n_{\text{pix}}^3}$ is the discretized domain of $V_i$ (and $V_j$).

Since we are working with 3D volumes, optimizing all coefficients $\lambda_{l,n,m}$ can be inefficient for large volumes. Instead, we propose working with a set of uniform 2D projections of the volumes and optimizing a single set of coefficients that is consistent across all projections:

$$\sum_j \left\| P_j \left(V_1(\mathbf{r})\right) - P_j \left(V_2(\mathbf{r} + \mathbf{g}(\mathbf{r}))\right) \right\|^2, \tag{27}$$

where $P_j$ denotes the projection operator along the $j$-th direction.

This new formulation significantly reduces the computational complexity of the problem, as it allows the use of the efficient optimization scheme proposed in [75]. By adjusting the number of projections, image resolution, and number of basis components, we can efficiently approximate the deformation field and derive a volume-to-volume distance with minimal compromise in accuracy. In our experiments we first down-sampled the volumes to box length of 64 (from 224), generated submission-specific masks using `xmipp_metadata.image_handler.ImageHandler.generateMask(..., iterations=50, boxsize=64, smoothStairEdges=False)` [76]. These yielded $\sim 3000 - 9000$ $(= 1000 - 3000 \times 3)$ spatial locations for the size of the domain of $\mathbf{g}(\mathbf{r})$. We used 20 $3D \to 2D$ projections to learn $120 \times 3$ Zernike coefficients for each spatial direction, and trained for 40 epochs. The computational implementation was parallelized across submissions, with a naive for loop over ground truth volumes.

#### 6.3 Rank based normalization of volume-to-volume distance matrix

We noticed that some ground truth or submitted volumes were on a higher or lower relative scale, or otherwise contained some bias that led to vertical or horizontal streaks of volume-to-volume distance matrices. This motivated us to rank normalize distance matrices, $\{d_{ij}\}$ to $\{R_{ij}\}$. We chose to (a) rank all submissions for one ground truth volume rather than (b) rank all ground truth volumes for a fixed for submitted volume, so that submitted volumes that were far from ground truth volumes would have a high rank. Concretely we used `scipy.stats.rankdata(..., method='average',` `axis=1`), which breaks ties according to the following criterion: "The average of the ranks that would have been assigned to all the tied values is assigned to each value".

#### 6.4 Results from different volume-to-volume distances: distance matrix, relative population, and objective value

The trends of the volume distribution objective (EMD or KL) depended on what volume-to-volume distance was assumed, as we had anticipated when launching the challenge. Typically the submissions that were the closest to the ground truth were the Average GT and Sampled GT, but not always when the population weights were optimized. We also compared results from the forward and reverse KL, and did not observe major differences (data not shown). In terms of the quantitative value of the EMD objective, lower volume-to-volume distances lead to lower EMD, as expected from the linear objective. Results with MSE, negative correlation, and resolution volume-to-volume distances had similar trends as the FSC distance and are omitted for brevity.

We found that the transport distances were brittle to comparing submissions against each other, especially Gromov-Wasserstein. Even when comparing Averaged GT to Sampled GT, which have a similar multiplicative and additive scale as well as sharpening, we found that the $\ell_2$ distance was more robust to the high frequency conformational heterogeneity of Sampled GT (Supplementary Fig. S4). The rotational residuals in the Procrustes-Wasserstein were very small (less than one degree), suggesting that either our alignment procedure was suitable, or are at least close to a local minimum. The Gromov-Wasserstein did not give the expected behavior for C2 symmetry, which would have given similar volume-to-volume distances for indices $i$ to $40 \pm j$, suggesting it has not reached a desirable optimum. We experimented with entropic regularization (`ot.gromov.entropic_gromov_wasserstein`) and our own local iterative solver with Frank-Wolfe (see Algorithm 2), but neither of these approached alleviated this issue. This could be due to a number of issues including differences of a local versus global solution to the objective, and also the value of the GW objective being dominated by 'high frequency' spatial displacements of mass.

We observed that Zernike3D could more smoothly measure the distance of a volume undergoing continuous conformational heterogeneity. As a forward mapping deformation distance it can displace volume intensity due to the conformational heterogeneity of thyroglobulin. We also observed negligible differences in the optimized relative population weights when the volume-to-volume distance was rank normalized. Supplementary Fig. S11 shows that for Zernike3D, the optimized population weights are similar under an KL and EMD objective (upper panels BC), but that trends among submissions from the objective value (lower panels BC) differ. We noticed that Chocolate Chip 2 was an outlier, with large values. We re-ran the volume to volume distance computational dozens of time and obtained similar 'vertical streaks' in the matrix, with maximum values ranging 4-6, although not always with identical submitted volumes, suggesting that the training was not reproducible, perhaps due to high noise in the submitted volumes, or undesired variability in the automatic mask generation routine.

As reference, we present results from the FSC distance (proportional to the area-under-the-FSC-curve). This volume-to-volume distance has also been employed in CryoBench [19], and was reasonably efficient to compute, while being fairly easy for us to interpret and debug, although some undesirable features have been pointed out in [29] (see their Fig. S4). In Supplementary Fig. S5A we show the volume-to-volume distance matrix for several representative submissions. Using these, we compare the optimal distribution (colored) to the submitted one (black) in the submission's space of volumes (Supplementary Fig. S5B). The conformational distribution (Supplementary Fig. S5C) of Averaged GT is close to the GT by both KL and EMD, as expected. Overall, this suggests we can reasonably compare smoothed volumes to the high frequency ground truth volumes with our validation pipeline, employing the FSC distance. In Supplementary Fig. S5C (lower) the optimized EMD only slightly decreases (light vs. dark bars for each submission), and in general the EMD values between submissions are much larger than this gap, even for re-submissions from the same group (numbered 1,2,...). This suggests that the FSC distance with the EMD, as a single optimizable scalar quantity between weighted ensembles of volumes, is (i) sensitive to the volume voxel intensities, (ii) fairly robust to their relative population, but (iii) can nevertheless be used to optimize the relative population weights in a reasonable manner.

#### 6.5 Volume distribution comparison: ground truth to submissions

Given (a) the distributions (weighted populations) of both ground truth and submitted volumes in their associated spaces, and (b) a volume-to-volume distance matrix between all pairs, we quantified the similarity between the distributions in two ways. The first was based on the Kullback–Leibler (KL) divergence, and the second was based on the earth mover's distance (EMD) from optimal transport theory. These objectives were chosen based not only on expected familiarity among the research community, but also because we could compute optimal submitted populations conveniently, which we think aids in the interpretability and validation of submitted population weights.

##### 6.5.1 KL divergence

We computed the KL over discrete patches of the heterogeneity space. We defined these patches as disjoint windows along the ground truth heterogeneity coordinate. Since the KL is defined pointwise, we chose a large enough window to smooth out the heterogeneity from the MD sampling.

In order to compute the KL between the ground truth and submitted relative population weights, the two vectors, which are each in their own simplex, need to be in a common space. We defined an association matrix *via* a 'hard' classification where each ground truth volume is associated with its nearest submitted volume. The association matrix was normalized such that all columns (submitted volumes) summed to 1. In the case where a submitted volume is an outlier and no ground truth volume is closet to it, it is associated with all ground truth volumes equally. The association matrix is thus:

$$A \in [0,1]^{m\times n}, \text{ s.t. } \forall j, \sum_i^m A_{ij} = 1. \tag{28}$$

After the association matrix is defined, the submitted population weight vector $q \in \Delta^{n-1}$ volumes to the ground truth space through the matrix multiplication

$$q' = Aq \in \Delta^{m-1}. \tag{29}$$

Finally, the KL is computed between $p$ and $q'$ both forward and reverse, respectively:

$$\text{KL}_f(p||q') = \Sigma_i p_i \log \frac{p_i}{q'_i} \ , \tag{30}$$

$$\text{KL}_r(q'||p) = \Sigma_i q'_i \log \frac{q'_i}{p_i} \ . \tag{31}$$

##### 6.5.2 Earth mover's distance (EMD)

Optimal transport theory naturally handles the problem of transforming distributions to each other (here $p$ and $q$), where there is a metric over their spaces (here the volume-to-volume distance). By solving the transport problem, we get the transport matrix that minimizes the cost of transporting probability between ground truth and submitted volumes, such that this cost is minimal. In other words:

$$\text{EMD}(p,q) = \text{EMD}(q,p) = \min_T \sum_{i,j} T_{ij} d_{ij}, \tag{32}$$

$$\text{s.t. } q_j = \sum_i T_{ij}, \text{ and} \tag{33}$$

$$p_i = \sum_j T_{ij} \ . \tag{34}$$

##### 6.5.3 Optimal submitted volume probabilities

Given the choice of the KL metric and EMD, we were conveniently able to optimize for $\{q_j\}_{j=1}^{n=80}$ given $\{t_i, p_i\}_{i=1}^{m=3861}$ and submitted $\{u_j\}_{j=1}^{n=80}$. This answers the question, given the ground truth volumes and relative population and also the participant's submitted volumes, what relative population of the participant's submitted volumes would have minimized the EMD or KL? This is an important question, since it is a target for cryo-EM inference methods focusing on source distribution estimation, which would estimate this directly from image data, as the participants in the challenge had to.

We found that despite differences in the objective and the association matrix $A$ versus the transport plan $T$, the optimal solution $q_*$ in both cases was very similar and also sparse (many elements were zero). In order to aid interpretability we typically visualized $q_*$ in plots where we smoothed with a running window, of default size 5 (except for Peanut Butter and Chocolate Chip 2, which seemed to be unordered, where no running window was assumed). Sparse solutions are expected for both objectives and there exist ways to regularize them in the objective itself, and we experimented with two types of regularization (data not shown): (i) negative entropy, and what we term (ii) self-distance. The latter promotes transport to be similar for submitted volumes that are similar to each other, and requires computing an $80 \times 80$ self-distance between submitted volumes, $d_{\text{self}}$, thus we have

$$\text{neg-entropy}(q) = \sum_i q_i \log q_i \tag{35}$$

$$\text{self-distance}(q) = \sum_i \sum_{jj'} \frac{(T_{ij} - T_{ij'})^2}{d_{\text{self},jj'}}. \tag{36}$$

Since $p$ is constant for the optimization objective, minimizing $\text{KL}_f$ is equivalent to the following objective

$$q'_* = \text{argmin}_{q \in \Delta^{n-1}} = -\Sigma_i p_i \log a_i^T q \tag{37}$$

where $a_i^T$ are the rows of $A$.

This KL objective can be solved with an iterative algorithm called multiplicative gradient descent, which keeps $q$ on the simplex for each gradient step [77]. We implemented this in Python, and used two termination criteria: (a) maximum iterations, with a default of $n_{\text{iter}} = 10^5$, and (b) multiplicative gradient whose elements each close to 0 or 1, up to an absolute tolerance of $10^{-4}$. The initial choice of $q_0$ was the uniform distribution.

Since the optimal transport objective is linear in $T_{ij}$ and since the constraints are linear in $T_{ij}$ and $q_j$, optimizing over both the matrix $T$ and the vector $q$ jointly reduces to a linear program (LP) [78]. We solved the system in the Python API of CVX [79, 80], which gave solutions with the ECOS solver with `max_iters` = 250, but otherwise default parameters. We formulated the problem with vectorized objective and constraints and it took several seconds to complete. For numerical stability we normalized the distance matrix by its absolute maximum, and multiplied it back in afterwards. This pre-normalization was critical when the volume-to-volume distance matrix had large values, and we could not overcome this through adjusting numerical tolerances. In the case of the un-regularized LP, this is equivalent to determining the closest submitted volume to each ground truth volume, and then normalizing the counts. This is because, when we drop the marginal constraint on the submitted probability and only have $\forall i, \quad p_i = \sum_j T_{ij}$, we can solve the optimization over $T$ row-wise by putting all the probability on $T_{ij*}$, where $j* = \arg\min_j d_{ij}$:

$$\min_T \sum_{i,j} T_{ij} d_{ij} = \sum_i \min_T \sum_j T_{ij} d_{ij} \tag{38}$$

For better statistics, we averaged EMD, KL and optimal probabilities over 30 random replicates. Each replicate used a random 90% subset of the ground truth volumes, uniformly sampled, and averaged the volume-to-volume distance matrix across segments of ground truth volumes ordered by the GT ordering (MD-PC1) in ground of ranging 1 to 50 (for a volume-to-volume distance matrix of $3861 \times 80$ to around $75 \times 80$). For some of the results shown we averaged the EMD or KL objective value over different segment sizes. In order to normalize EMD we divide by the segment size (effectively using the mean of the volume-to-volume matrix in the neighborhood), while the KL requires no normalization since the columns of A always sum to 1, regardless of the number of rows.

Note that as part of the validation we randomly subset ground truth volumes, average the volume-to-volume distance over regions of the GT ordering. The solutions are sparse (some optimal populations are driven to zero), and for visualization purposes we averaged the optimized population over a small window of 5 nearest neighbors where an ordering way apparent in the submission, and did not average over neighbors for submissions with seemingly unordered volumes. Note that there was no averaging for the actual optimal value used in computing the optimized EMD in Fig. 3C, which is therefore order independent.

#### 6.6 Image comparison metrics: cross-correlation and ensemble likelihood

In this challenge, we developed metrics to compare the submitted volumes and population estimates to the ground truth. However, we also aim to establish metrics that evaluate how well the submissions capture the underlying image space, i.e., metrics that can be applied even in the absence of ground truth. To compute the cross-correlation as well as the image-to-volume likelihood, $p(y_i|V_m)$, we use cryoLike [33]. This package has been developed to calculate the cross-correlation and likelihood of an individual image to a given volume using a Fourier-Bessel basis and a white noise model, exploiting the degeneracy of in-plane rotations in this basis. The algorithm searches on a grid for the center of the template particle, and integrates the image-to-structure likelihood over a grid of possible poses over the orientation sphere. To choose the hyperparameters for the cryoLike algorithm, we performed several trials, sweeping across different orientation searches and centroid positions and prior distributions. We settled on a center search of 1 grid point per pixel, 2370 templates around the orientation sphere, and 256 unique in-plane rotations. This resulted in roughly 600,000 unique poses. To limit computational costs, we limit likelihood comparisons to 8.6 Å with a low-pass filter in Fourier space. We limited the center search extent to within 15 pixels of the particle center. Once the whole orientation sphere has been searched, we record the maximum cross-correlation between the image and the projected template. This is reported in Fig. 2C.

To compute the ensemble-to-image likelihood, we build on the Bayesian inference pipelines that estimate a likelihood given a set of volumes $\{V_m\}$ with population weights $\{\alpha_m\}$ and an image set $Y = \{y_i\}$ [32, 66]. Hence, for each particle image the likelihood of a user's submission can be calculated as

$$p(y_i|\{V_m, \alpha_m\}) = \sum_m p(y_i|V_m)\alpha_m \ . \tag{39}$$

We note that we are not performing any optimization over the weights but using the participant's submitted values, and $p(y_i|V_m)$ is computed using cryoLike as described above.

To avoid comparing ensembles with different numbers of volumes (submissions have 80 while GT has 3861), for the simulated data we only analyzed a subset of 1224 images coming from 80 GT volumes (Sampled GT). For the experimental dataset, we selected only 2000 images for computational simplicity. Note that the computational expense is about 1-2 GPU hours per submission (80 volumes) for 2000 images (downsampled to 112 pixels).

### Supplementary Material

#### List of Supplemental Movies.

1. “Supplemental Movie 1”: *Supplemental_Movie_Experimental_Motions.mp4*, a movie of the submitted volume series for the experimental dataset colored as in Figure 2A.
2. “Supplemental Movie 2”: *Supplemental_Movie_GT_latent.mp4*, a movie of the Sampled Ground Truth volumes along the ground truth latent.

**Table 1**: Pre-processing summary for set 1, round 1.

| Submission | **Box Size** | **Voxel Size** | **Initial Manual Alignment** | **Flip** | **#Volumes** |
|---|---|---|---|---|---|
| Black Raspberry 1 | 224 | 2.146 | Identity | No | 80 |
| Chocolate 1 | 224 | 2.146 | Identity | No | 76 |
| Chocolate Chip 1 | 224 | 2.146 | Identity | No | 80 |
| Cookie Dough 1 | 224 | 2.146 | Identity | No | 80 |
| Mango 1 | 224 | 2.146 | Identity | No | 80 |
| Neapolitan 1 | 224 | 2.146 | Identity | No | 80 |
| Peanut Butter 1 | 224 | 2.146 | Identity | No | 80 |
| Rocky Road 1 | 284 | 1.073 | Identity | No | 80 |
| Salted Caramel 1 | 224 | 2.146 | Identity | No | 80 |
| Vanilla 1 | 224 | 2.146 | $y = -90^\circ$ | No | 80 |

**Table 2**: Pre-processing summary for set 1, round 2.

| Submission | **Box Size** | **Voxel Size** | **Initial Manual Alignment** | **Flip** | **#Volumes** |
|---|---|---|---|---|---|
| Black Raspberry 2 | 224 | 2.146 | Identity | No | 80 |
| Chocolate 2 | 224 | 2.146 | Identity | No | 80 |
| Neapolitan 2 | 224 | 2.146 | Identity | No | 80 |
| Peanut Butter | 224 | 2.146 | Identity | No | 80 |
| Piña Colada 2 | 224 | 2.146 | $z = -90^\circ$ | No | 80 |
| Salted Caramel 2 | 224 | 2.146 | Identity | No | 80 |
| Salted Caramel 3 | 224 | 2.146 | Identity | No | 80 |
| Vanilla 2 | 224 | 2.146 | Identity | No | 80 |

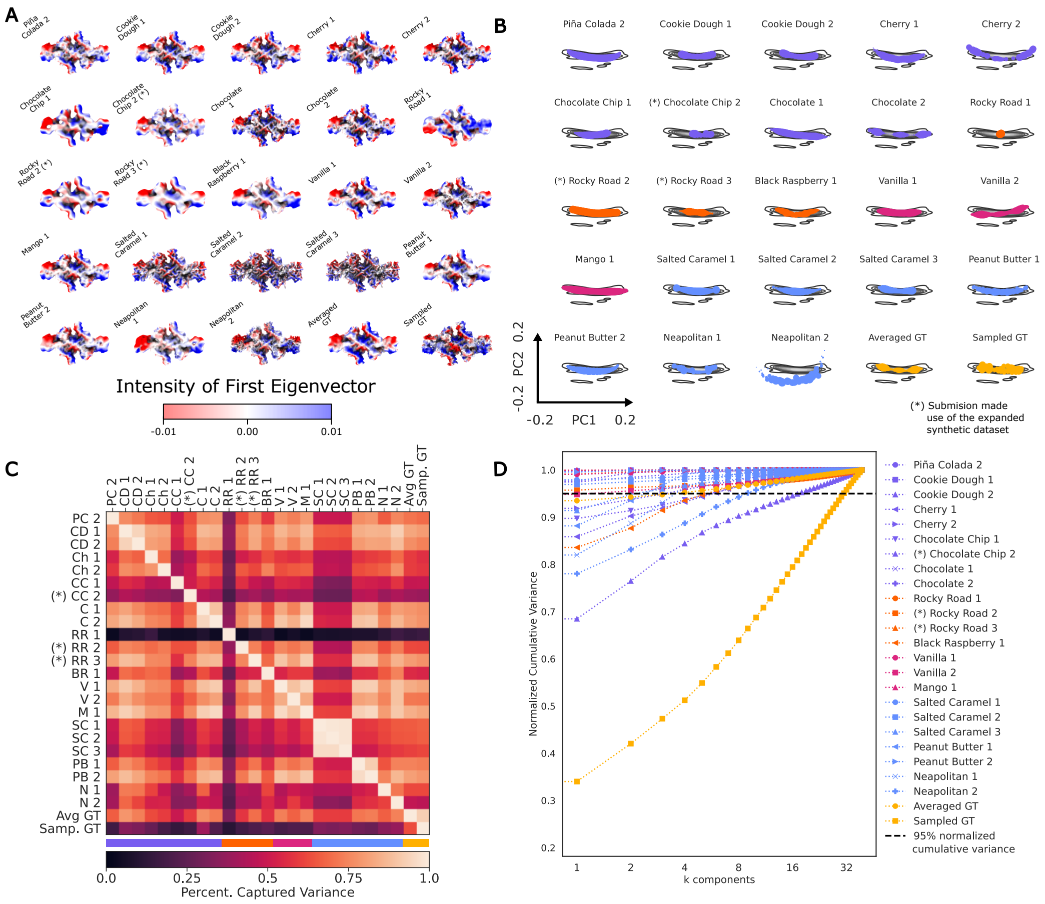


**Fig. S1**: PCA analysis of submissions from the synthetic set. A) The 40th volume of each submission to the synthetic dataset colored by the local intensity of the first eigenvector for that submission as determined by SVD. B) shows the common embedding computed from the submissions. Most submissions recover the same motion in the first principal component and are well aligned with the motions for both ground truth mock submissions. In C) we present the PCV between all submissions. The rows correspond to the reference subspace, (see Section 6.1 for details), and the colored bar on the top shows the category assigned to each group. We observe that methods in the same category tend to form clusters (high PCV regions close to the diagonal), and that methods with a higher resolution are not captured as well as low resolution methods (see columns and rows associated to Sampled GT versus Averaged GT). D) Shows the cumulative normalized variance, defined for each submission as $\sum_i^k \sigma_i^2 / \sum_i \sigma_i^2$, where $\sigma_i$ is the i-th singular value of a given submission.

**Table 3**: Pre-processing summary for set 2, round 1.

| Submission | **Box Size** | **Voxel Size** | **Initial Manual Alignment** | **Flip** | **#Volumes** |
|---|---|---|---|---|---|
| Black Raspberry 1 | 224 | 2.146 | Identity | No | 80 |
| Cherry 1 | 224 | 2.146 | Identity | No | 80 |
| Chocolate 1 | 224 | 2.146 | Identity | No | 79 |
| Chocolate Chip 1 | 168 | 2.146 | Identity | No | 80 |
| Cookie Dough 1 | 128 | 2.146 | $z = -60°$ | No | 80 |
| Mango 1 | 128 | 2.146 | Identity | No | 80 |
| Neapolitan 1 | 288 | 1.073 | $x = 180°$ | No | 80 |
| Peanut Butter 1 | 224 | 2.146 | Identity | No | 80 |
| Rocky Road 1 | 284 | 1.073 | Identity | No | 80 |
| Salted Caramel 1 | 144 | 2.146 | Identity | No | 80 |
| Vanilla 1 | 224 | 2.146 | $y = -90°$ | No | 80 |

**Table 4**: Pre-processing summary for set 2, round 2.

| Submission | **Box Size** | **Voxel Size** | **Initial Manual Alignment** | **Flip** | **#Volumes** |
|---|---|---|---|---|---|
| Cherry 2 | 224 | 2.146 | Identity | No | 80 |
| Chocolate 2 | 220 | 2.146 | Identity | No | 80 |
| Chocolate Chip 2 | 288 | 1.67 | $z = 90°$ | No | 80 |
| Cookie Dough 2 | 256 | 1.073 | $(z, x) = (60°, 180°)$ | Yes | 80 |
| Neapolitan 2 | 288 | 1.073 | Identity | Yes | 80 |
| Peanut Butter 2 | 224 | 2.146 | Identity | No | 80 |
| Piña Colada 2 | 224 | 2.146 | Identity | No | 80 |
| Rocky Road 2 | 448 | 1.073 | $z = 90°$ | No | 80 |
| Rocky Road 3 | 448 | 1.073 | $z = 90°$ | No | 80 |
| Salted Caramel 2 | 288 | 1.073 | Identity | No | 80 |
| Salted Caramel 3 | 288 | 1.073 | Identity | No | 80 |
| Vanilla 2 | 144 | 2.146 | Identity | Yes | 80 |

| Cistem Simulator Parameters | Value |
|---|---|
| Ice Thickness Å | 450 |
| Total dose $e^-/Å^2$ | 30 |
| Propagation distance Å | 5 |
| Number of frames | 30 |
| Per-atom b-factor scaling | 1 (default) |
| Per-atom b-factor constant | 5 (default) |
| Pre-exposure dose $e/Å^2$ | 1 |
| Image size px | 288 × 288, or 448 × 448 |
| Pixel size Å | 1.073 |
| Exposure Filtering (developer option) | Yes |

**Table 5**: Input for the cisTEM simulator during the generation of the synthetic dataset.

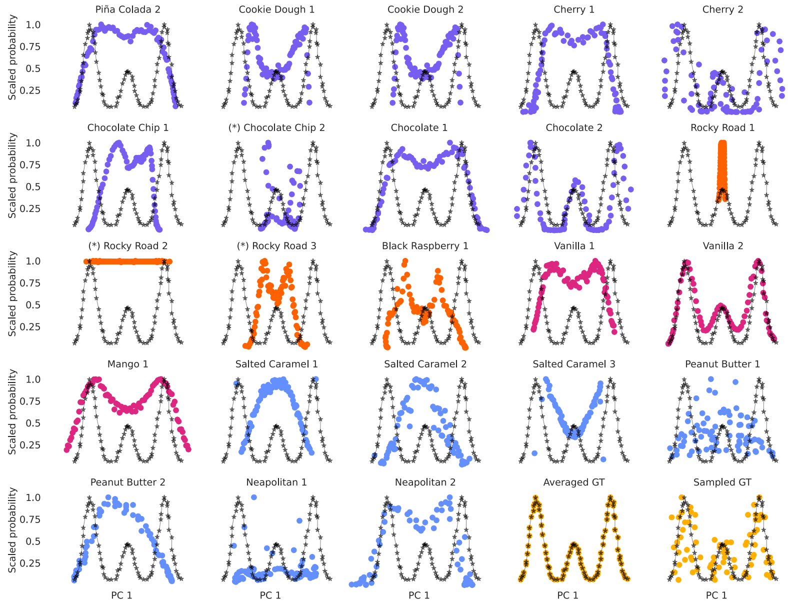


**Fig. S2**: Submitted populations projected onto the first principal component of the Sampled GT (x-axis and see Section 1.4). The black dots correspond to the populations of the Averaged GT. The coverage of the x-axis indicates that most submissions manage to recover the motion in the first principal component. However, most methods fail at reporting the three modes of the GT distribution.

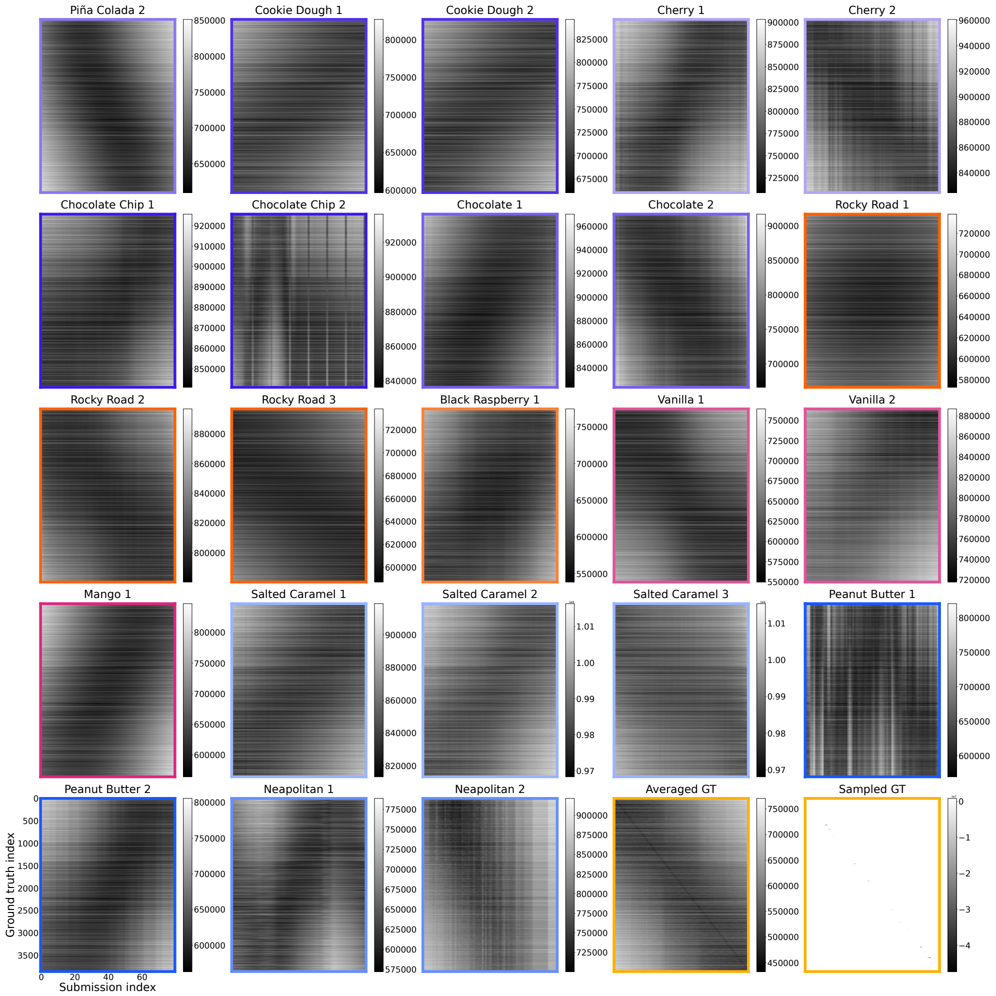


**Fig. S3**: Volume-to-volume distance matrices for all submitted volumes for the simulated set to the 3861 GT Volumes under the BioEM3D metric. Sampled GT has low values near the diagonal, which can be faintly seen in the rendering of the figure due to the large number of rows.

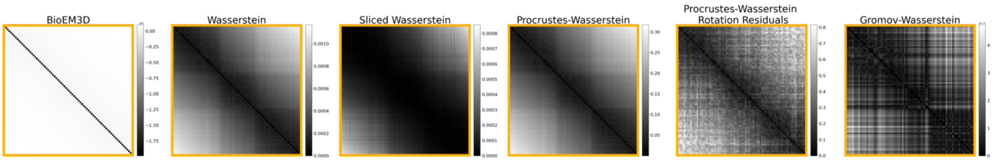


A: Averaged GT vs. itself: volume-to-volume distances

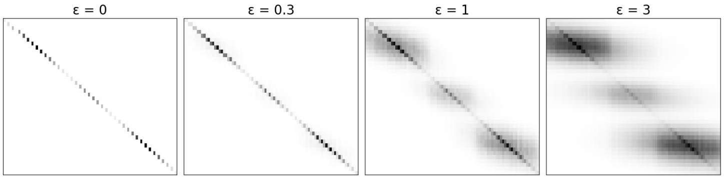


B: Averaged GT vs. itself: transport plans

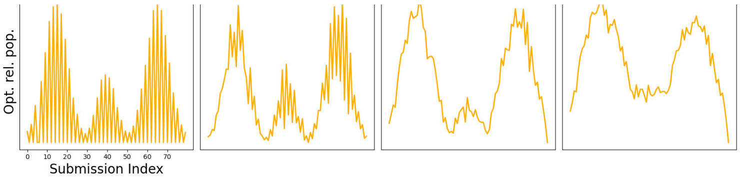


C: Averaged GT vs. itself: relative populations

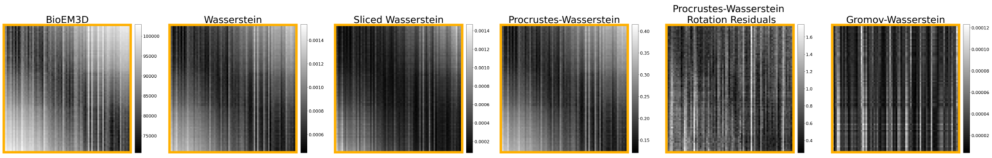


D: Averaged GT vs. Sampled GT: volume-to-volume distances

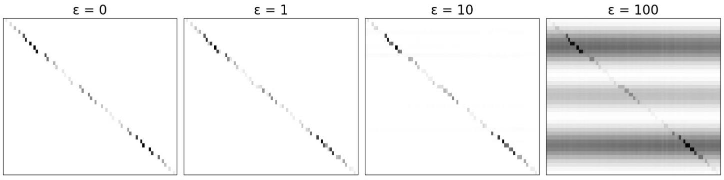


E: Averaged GT vs. Sampled GT: transport plans

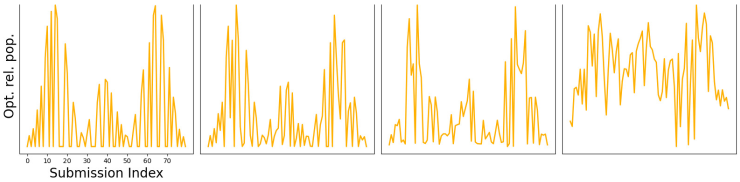


F: Averaged GT vs. Sampled GT: relative populations

**Fig. S4**: **(A,D): BioEM3D compared with transport distances**. We computed volume-to-volume transport distances of Averaged GT volumes to themselves (A) and to Sampled GT (D). **(B,C,E,F): Conformational distribution: EMD with entropic regularization.** We compared the EMD under the BioEM3D volume-to-volume distance regularized with the neg-entropy term (Eq. 35). The neg-entropy was weighted by $\epsilon$ relative to the EMD term, where the volume-to-volume distance matrix has been divided by the maximum. The GT was averaged in a GT ordering (MD-PC1) window of 95, to make an averaged cost matrix of $40 \times 80$, which can only have a maximum of 40 non-zero entries with no regularization (cf. commentary on Eq. (38)). The transport plans (B,E) and the optimized relative population (i.e. the marginal of the transport plan) (C,F) are shown. For Sampled GT, even at large $\epsilon = 100$, the submitted volumes (columns) with large offsets (i.e. vertical streaks in the BioEM3D volume-to-volume matrix) are down-weighted in the transport plan (corresponding vertical streaks).

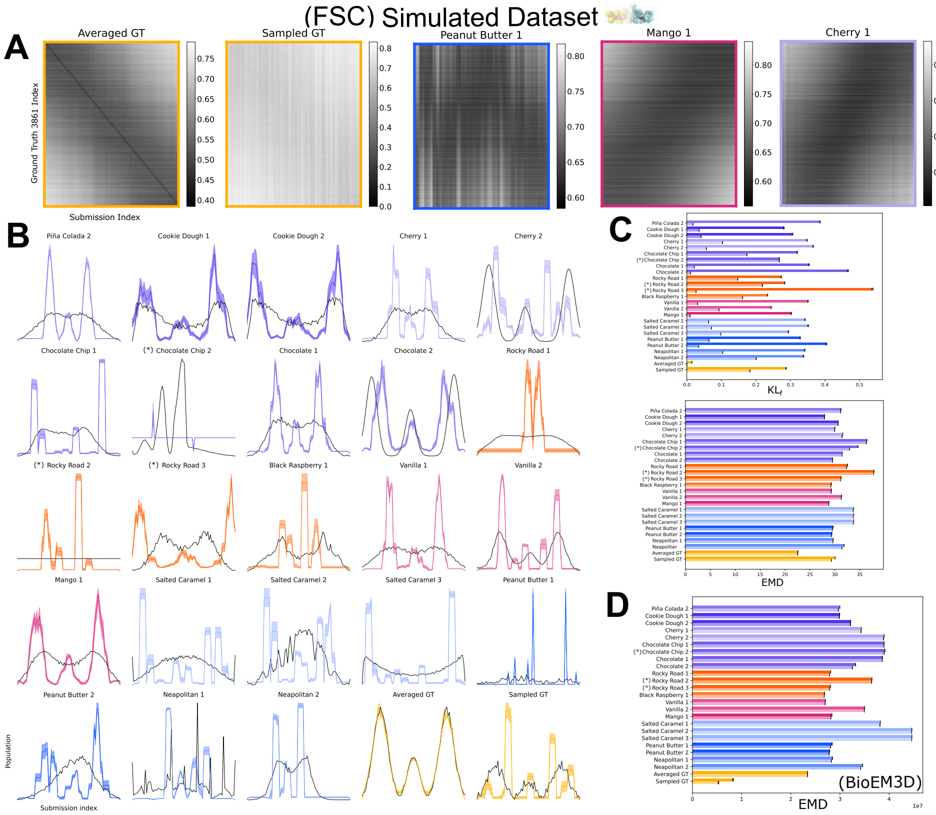


**Fig. S5**: **A.** Volume-to-volume FSC distance, Eq. (21), for select submissions against 3861 GT volumes. **B.** Optimal relative population *via* KL (in color, Eq. (33)) vs. original submitted relative population weights (black). Replicates subsample ∼ 90% (3474 of 3861) GT volumes, 30 times. **C.** Optimization metrics under KL (upper) and **E**MD (lower) using the FSC distance. **D.** Optimization under metrics for EMD using the BioEM3D (sic) distance. The error bars are from averaging the objective values when GT ordering (MD-PC1) neighborhoods of 40,45,50 were used.

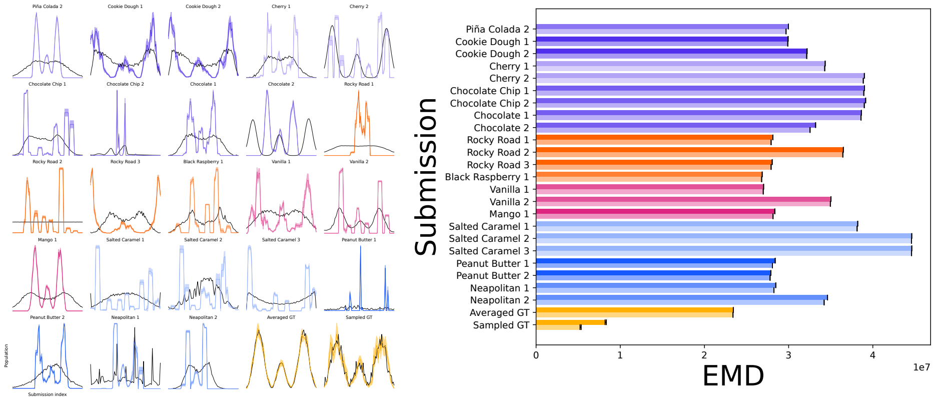


A: Relative Population and EMD objective value

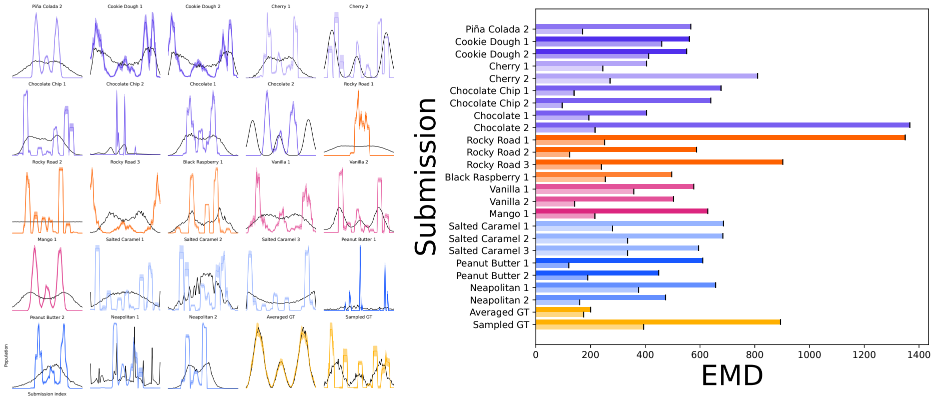


B: Relative Population and rank normalized EMD objective value

**Fig. S6**: **The optimized relative population is robust to rank normalization**. **A.** The optimized relative populations (left) and the EMD metric (right) for the BioEM3D distance matrix *without* rank normalization. **B.** The optimized relative populations (left) and the EMD metric (right) for the BioEM3D distance matrix *with* rank normalization (section 6.3). The relative populations do not change with rank normalization, with the exception of Sampled GT. This may be due to the low values along the diagonal of the distance matrix for this volume series when compared to the full GT (Figure 3A).

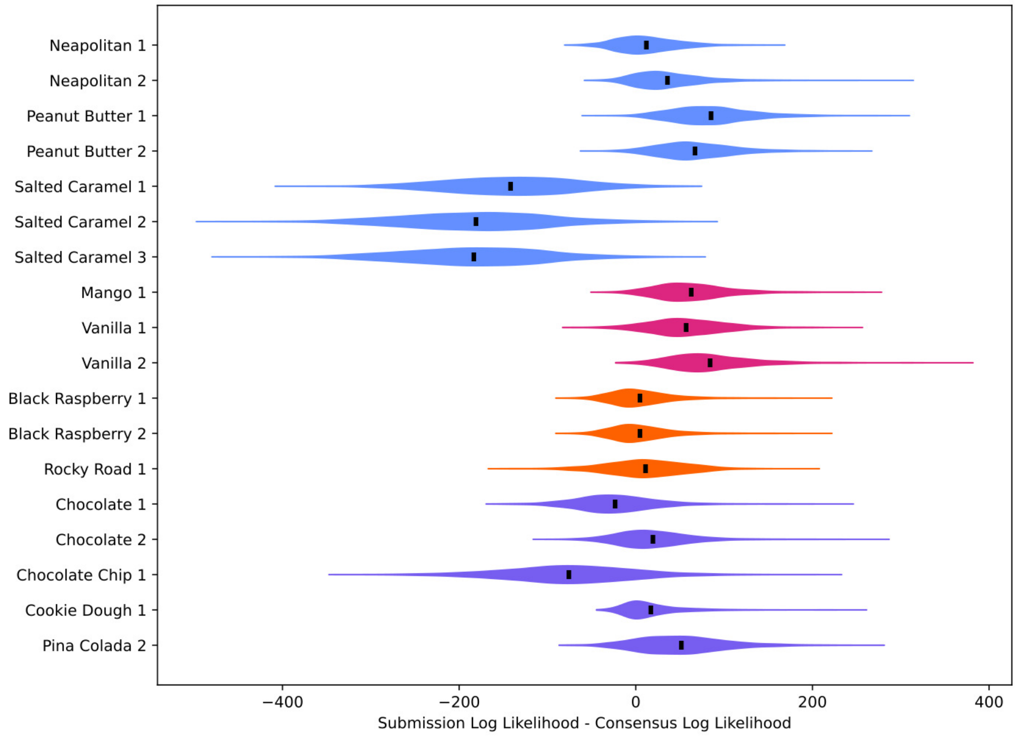


**Fig. S7**: Ensemble-to-image log likelihood results calculated for a subset of 2000 particles from the experimental dataset, calculated with the cryoLike likelihood[33].

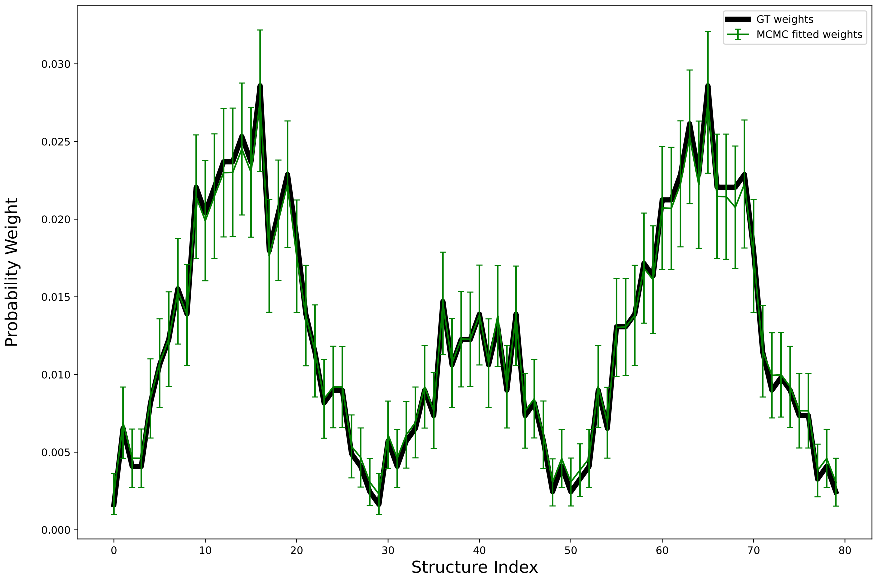


**Fig. S8**: **Correct Recovery of Relative Weights using MCMC and CryoLike**. In the simulated dataset, we can take the 80 volumes from the Sampled Ground Truth submission and calculate the structure-image likelihood for the 1224 images which correspond to the 80 volumes in the submission. When the relative weights of these volumes are estimated using Markov Chain Monte Carlo, we recover the correct weights of all 80 volumes.

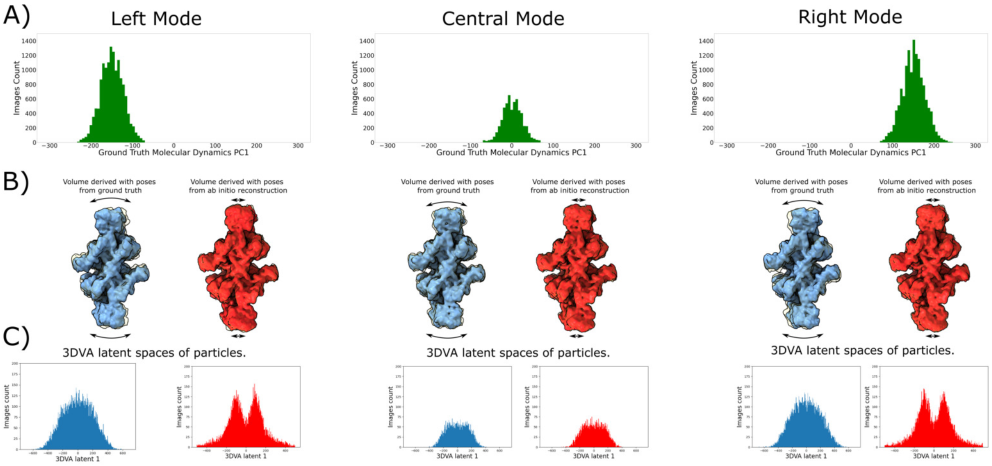


**Fig. S9**: **A.** Particles are selected according to their ground truth latent coordinate from the molecular dynamics simulation. **B.** Reconstructions are produced from either the ground truth poses or poses derived from ab initio reconstruction. The volumes produced with poses from ab initio reconstruction have a smaller range of motion than those derived using the ground truth poses. **C.** Analysis using 3DVA indicates that the reconstructions from the ground truth poses contain a uni-modal structural ensemble, while the reconstructions from ab initio reconstruction contain a bimodal structural ensemble.

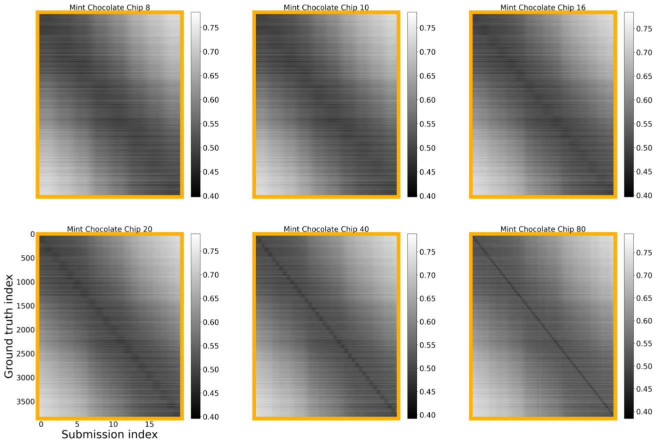


A: Volume-to-volume distance (FSC)

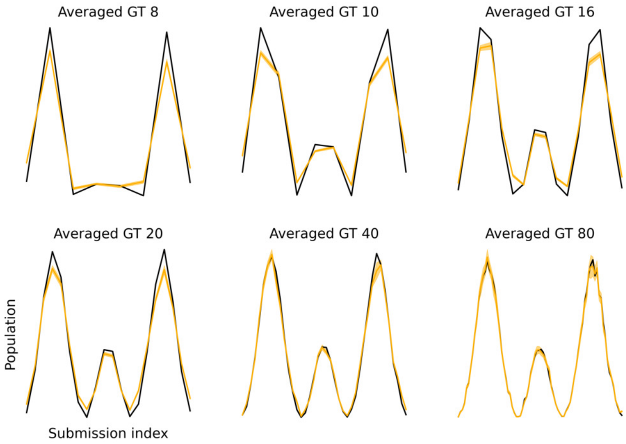


B: Optimal probability

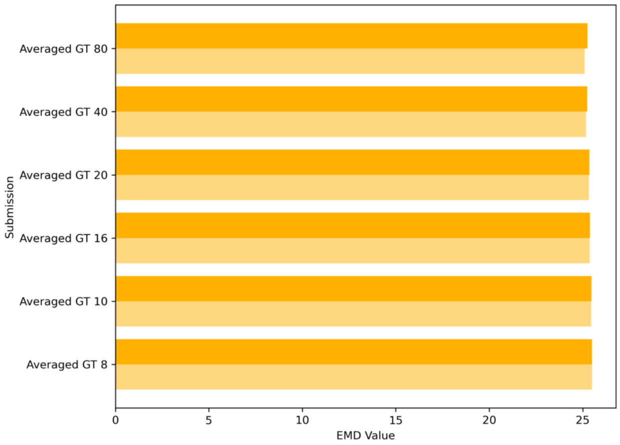


C: Distribution distance

**Fig. S10**: We averaged together ground truth subsets of various sizes (Averaged GT 80, 40, 20, 16, 10, 8) to make smoother volumes of smaller population size. **A.** Volume-to-volume FSC distance. **B.** Optimal probability *via* EMD. The ground truth population weights were averaged accordingly and are shown in black. The optimal probability is shown with no neighbor-averaging (i.e., window size = 1; see Section 3.4) **C.** EMD between ground truth and mock submission, with (lighter) and without (darker) optimal probabilities.

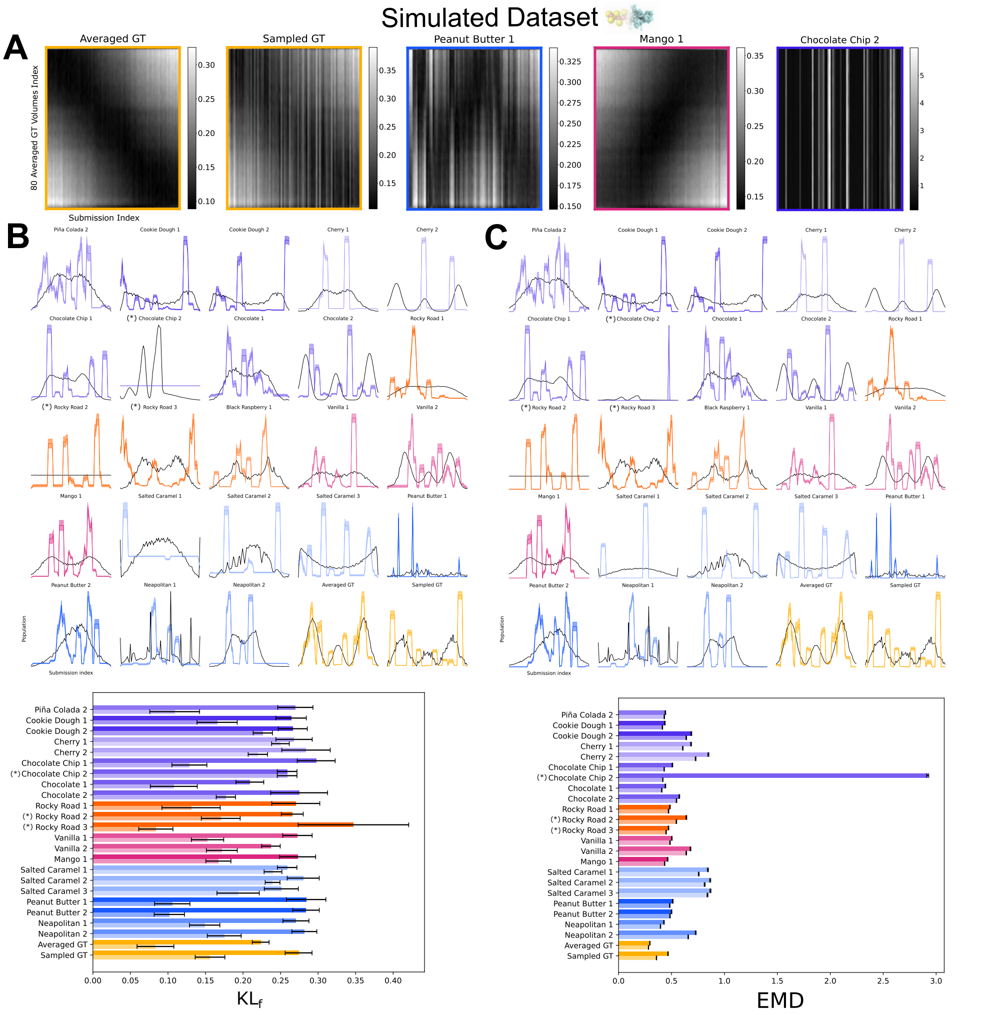


**Fig. S11**: **A.** Volume-to-volume Zernike3D distance, Eq. (26), for select submissions against Averaged GT. Optimal relative population (in color, Eq. (33)) *via* KL (**B.**) and EMD (**C.**), and original submitted relative population weights (black). Replicates subsample 90% (72 of 80) averaged GT volumes, 30 times. Bottom panels compare KL and EMD optimization metrics. The error bars are from averaging the objective values when (Averaged) GT MD-PC1 neighbourhoods of 1-5 were used.
